# Effects of Shift Work on the Carotid Artery and Cerebral Blood Flow of Spontaneously Hypertensive Rats and Wistar-Kyoto Rats

**DOI:** 10.1101/740068

**Authors:** YunLei Wang, Tong Zhang, YuGe Zhang, Yan Yu, Fan Bai, HaoJie Zhang, YaFei Chi, Shan Gao

## Abstract

**Objective:** The objective was to investigate the effects of shift-work (SW) on the carotid arteries.

**Methods:** This study used two inverted photoperiods (inverted light:dark [ILD]16:8 and ILD12:12) to create the SW model. We recorded the rhythm and performed serological tests, carotid ultrasound, magnetic resonance imaging, and carotid biopsy.

**Results:** SW induced elevated blood pressure and increased angiotensin-II, apolipoprotein E, blood glucose, and triglycerides. SW increased the carotid intima-media thickness. SW led to the development of carotid arterial thrombosis, reduced cerebral blood flow, and increased the number of collagen fibers, expression of angiotensin receptor and low-density lipoprotein receptor in the carotid arteries. SW decreased 3-hydroxy-3-methylglutaryl-CoA reductase and nitric oxide. SW induced the atherosclerotic plaque in the aorta. Multiple results of SHR were worse than WKY rats.

**Conclusion:** SW can induce metabolic disorders and elevated blood pressure. SW can cause intima-media thickening of the carotid artery and aorta atherosclerosis. SW may result in carotid arterial thrombosis and affect cerebral blood flow. Hypertension can aggravate the adverse effects of SW.

## Introduction

Despite the remarkable advancements in the field of medicine, the increase in the incidence of stroke has not yet been effectively managed. According to an epidemiological sample survey on cerebrovascular disease, the incidence of ischemic stroke in China reached 0.2468%, and showed a trend of sustained increase with decreasing patient age[^1^]. In the past few decades, studies identified important risk factors related to stroke and its pathogenesis (i.e., blood sugar metabolism, dyslipidemia, hypertension, smoking, certain drug intake, and mental factors). Effective control of these risk factors will significantly reduce the incidence of stroke; however, this requires determination of their causes. Numerous relevant studies have been performed. However, the reasons for the increasing incidence of stroke in some countries in parallel with a decreasing age of patients affected by stroke, remain to be determined.

Most organisms have approximately 24-h circadian rhythms, which is a free-running endogenous rhythm[^2^]. However, owing to the effects of the Earth’s rotation and Sun’s illumination, most organisms form a 24-h rhythm that is synchronized to the external environmental factors, which is termed diurnal rhythm[^3^]. Through the entrainment of the “Zeitgeber” (ZT) [^4^], the organism maintains the endogenous circadian timing system in sync in response to changes in the external environment [^5^]. This ability maintains the body in an orderly and healthy state. When the biological clock is no longer consistent with the rhythm of the exogenous environment, such as in the case of shift work (SW), the creature may suffer a variety of negative effects and diseases[^6, 7^]. In recent years, owing to the rapid development of the economy and obvious acceleration of the pace of life, the number of shift workers has increased significantly[^8^]. Strikingly, in developed countries, approximately 20% of the population is involved in SW[^9^]. Similarly, in the United States of America, approximately 40% of workers are involved in SW[^10^]. In general, the greatest hazard of SW is that the endogenous rhythm does not result in adaptive adjustments in response to changes in the activity rhythm induced by exogenous factors[^11, 12^]. This means that the physiological rhythm of the organism cannot be changed with the activity rhythm after the inverted, which inevitably leads to the development of a series of diseases. Considering the current trends of the decreasing age of patients with stroke and continuous increase in the incidence of stroke, we strongly suspect that SW may play a major role in this process. Some studies have suggested a correlation between SW and the risk factors of cerebrovascular disease. For example, a clinical trial confirmed that SW induced increased variability in arterial blood pressure [^13^]. A retrospective study conducted by Gamaldo *et al*. also reported that the incidence of coronary heart disease was higher in patients with hypertension than in normotensive subjects [^14^]. A few retrospective reports suggested that SW may be correlated with metabolic disorders[^15^], coronary arterial atherosclerosis [^16^], and ischemic stroke [^17^]. Animal studies demonstrated that SW increased the blood pressure[^18^] and induced metabolic disorders[^19, 20^]. These outstanding studies suggested that SW may induce and even aggravate the risk factors of atherosclerosis, (e.g., hypertension and metabolic disorders). Unfortunately, there is no direct study investigating the effect of SW on the carotid arteries.

Another fact that needs people’s attention is that the “eat more, less active” lifestyle has led to an increase in the prevalence of hypertension. In China, a 2015 epidemiological survey reported that 27.9% of adults had hypertension[^21^]. Unfortunately, currently, there is a lack of experiments on the impact of SW on patients with hypertension. Therefore, this study used spontaneously hypertensive rats (SHR) and Wistar-Kyoto (WKY) rats as the homologous control group of SHR, to investigate the different effects of SW on the carotid arteries and cerebral blood flow (CBF).

Common methods for the regulation of the activity rhythm include photic entrainment [^22^] and nonphotic entrainment[^23^]. Light illumination transmits light information to the suprachiasmatic nucleus only through the retina and optic nerve. Therefore, the suprachiasmatic nucleus coordinates the timing of the tissue rhythms with each other and with the environmental light:dark (LD) cycle[^24, 25^]. Thus, different photoperiods are the most common regulator of the activity rhythm in all organisms. [^26, 27^] For the simulation of the adverse effects of SW (LD cycles) caused by conventional illumination in daily living [^28, 29^], we interfered with the activity rhythm of rats using ordinary fluorescent lamps (50–100 W) at typical illumination intensities (150–200 lux) to induce different LD cycles. At present, the two relatively mature and common SW regulation models in animal experiments are the following: [^30–32^] (1) inverted LD (ILD) activity, (represented by the ILD12:12 model in this study); and (2) long-term illumination with artificial light sources at night that causes imbalance of the LD time ratio and inverted or LD activity (represented by the ILD16:8 model in this study) (see Methods section for specific model establishment methods).

## Experimental results

### 1. ClockLab behavioral analysis

Period: Compared with LD12:12, ILD12:12 and ILD16:8 significantly prolonged the time of the period (F=36.651, *p*<0.001), with the ILD16:8 period being significantly longer than the ILD12:12 period (*p*<0.05) (Figure 2A). The difference in the time of the period between the SHR and WKY rats under the same LD conditions was not significant (F=0. 393, *p*>0.05).

**Figure 1:**
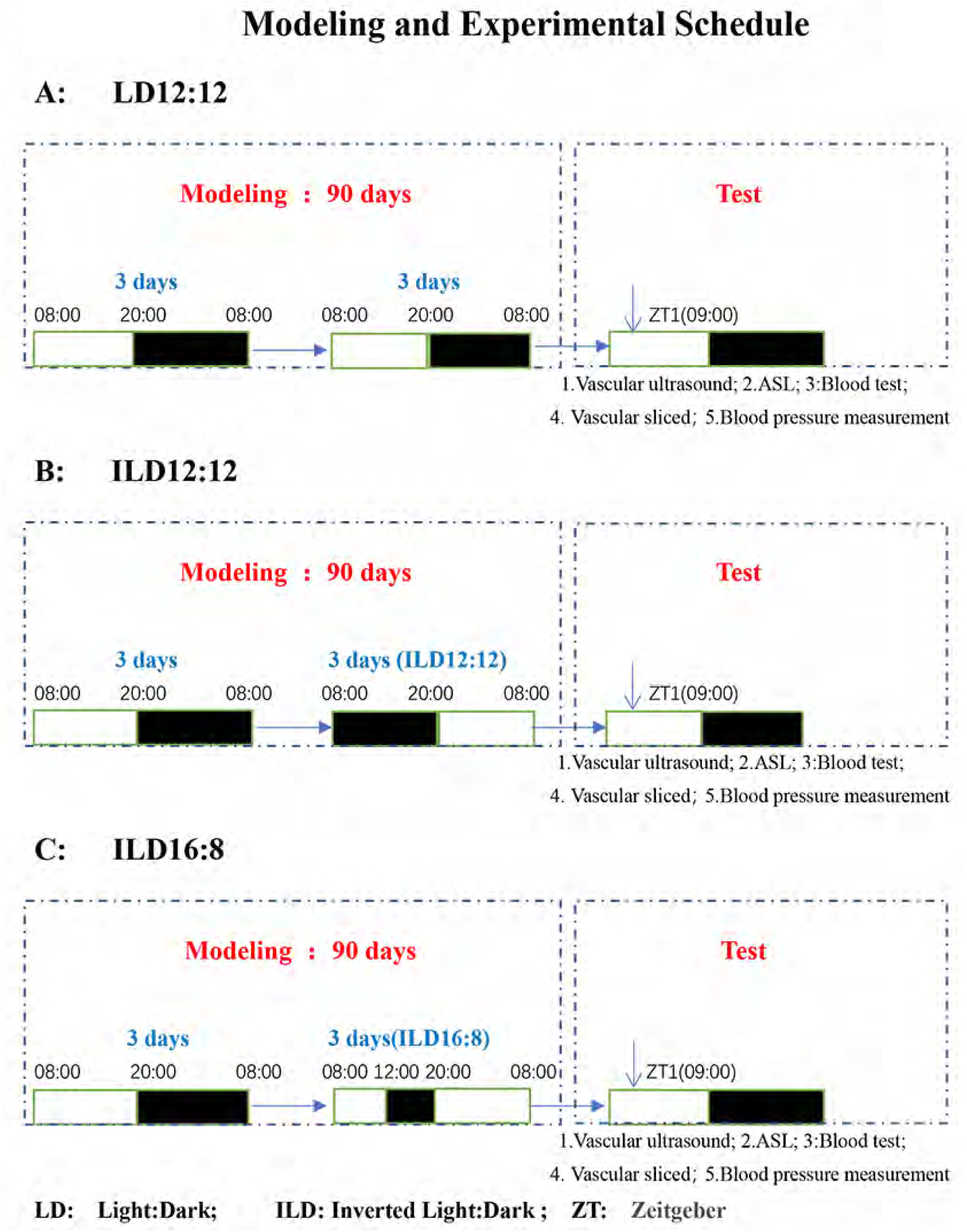
The three different photoperiods in this study established with different LD cycles are shown. Figure 1A, 1B, and 1C show the modeling and test time of the LD12:12, ILD12:12, and ILD16:8 groups, respectively. The model establishment period was 90 days. The measurement period was on days 91-93. Measurements in the three groups of rats were started at ZT1 (09:00).

**Figure 2:**
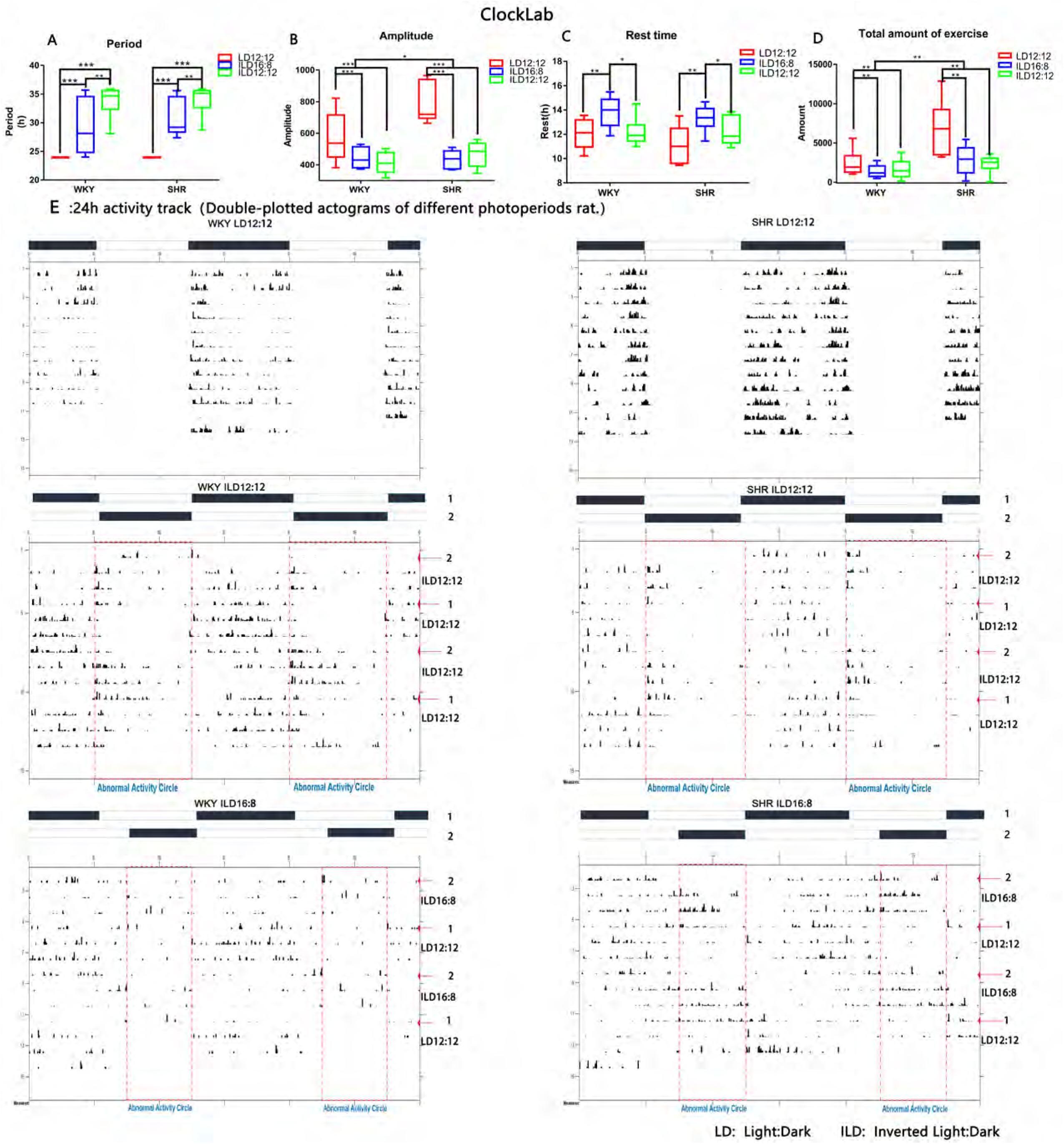
A-D, rhythm period, rhythm amplitude, rest time, and total exercise data of the six groups of rats. * *p* < 0.05, ** *p* < 0.01, *** *p* < 0.001. A two-factor design was used to compare the six groups of rats. The *post-hoc* Bonferroni was used for comparisons between groups. E, Double-plotted actograms of different photoperiods rat.

Rhythm amplitude: Compared with LD12:12, ILD12:12 and ILD16:8 significantly reduced the rhythm amplitude (F=22.121, *p*<0.001); however, the difference between the ILD16:8 and ILD12:12 was not significant (*p*>0.05) (Figure 2B). Under the same LD, the rhythm amplitude of the SHR was markedly higher than that observed in the WKY rats (F=6.724, *p*<0.05).

Rest time: The rest time in the ILD16:8 group was significantly longer than those reported in the LD12:12 (*p*<0.01) and ILD12:12 (*p*<0.05) groups (Figure 2C). However, there was no significant difference in the rest time between the ILD12:12 and LD12:12 groups (*p*>0.05). Under the same LD conditions, there was no significant difference in the rest time between the SHR and WKY rats (F=3.582, *p*>0.05).

Number of running wheel revolutions: Compared with LD12:12, ILD12:12 and ILD16:8 significantly reduced the number of running wheel revolutions (F=7.228, *p*<0.01) (Figure 2D). However, there was no significant difference in the number of revolutions between the ILD16:8 and ILD12:12 groups (*p*>0.05). However, under the same LD conditions, the number of running wheel revolutions in the SHR was significantly higher than that observed in the WKY rats (F=11.640, *p*<0.01).

Compared with LD12:12, ILD12:12 and ILD16:8 significantly prolonged the time of the period, indicating that SW caused the activity rhythm to be out of sync with the activity rhythm during the normal photoperiod. Furthermore, SW resulted in decreased activity rhythm amplitude and decreased exercise. The rest time of the ILD16:8 group was significantly longer than those reported in the LD12:12 and ILD12:12 groups.

### 2. Analysis of blood pressure

Figures 3A–3C represent the statistical results for the mean arterial pressure, systolic blood pressure, and diastolic blood pressure of rats prior to the experiment. The results showed that the differences in the mean blood pressure (F=486.432, *p*<0.001), systolic blood pressure (F=307.775, *p*<0.001), and diastolic blood pressure (F=384.869, *p*<0.001) between the SHR and WKY rats were significant prior to the experiment. However, the differences in the mean blood pressure, systolic blood pressure, and diastolic blood pressure among the three photoperiod groups of SHR and WKY rats were not significant (*p*>0.05).

**Figure 3:**
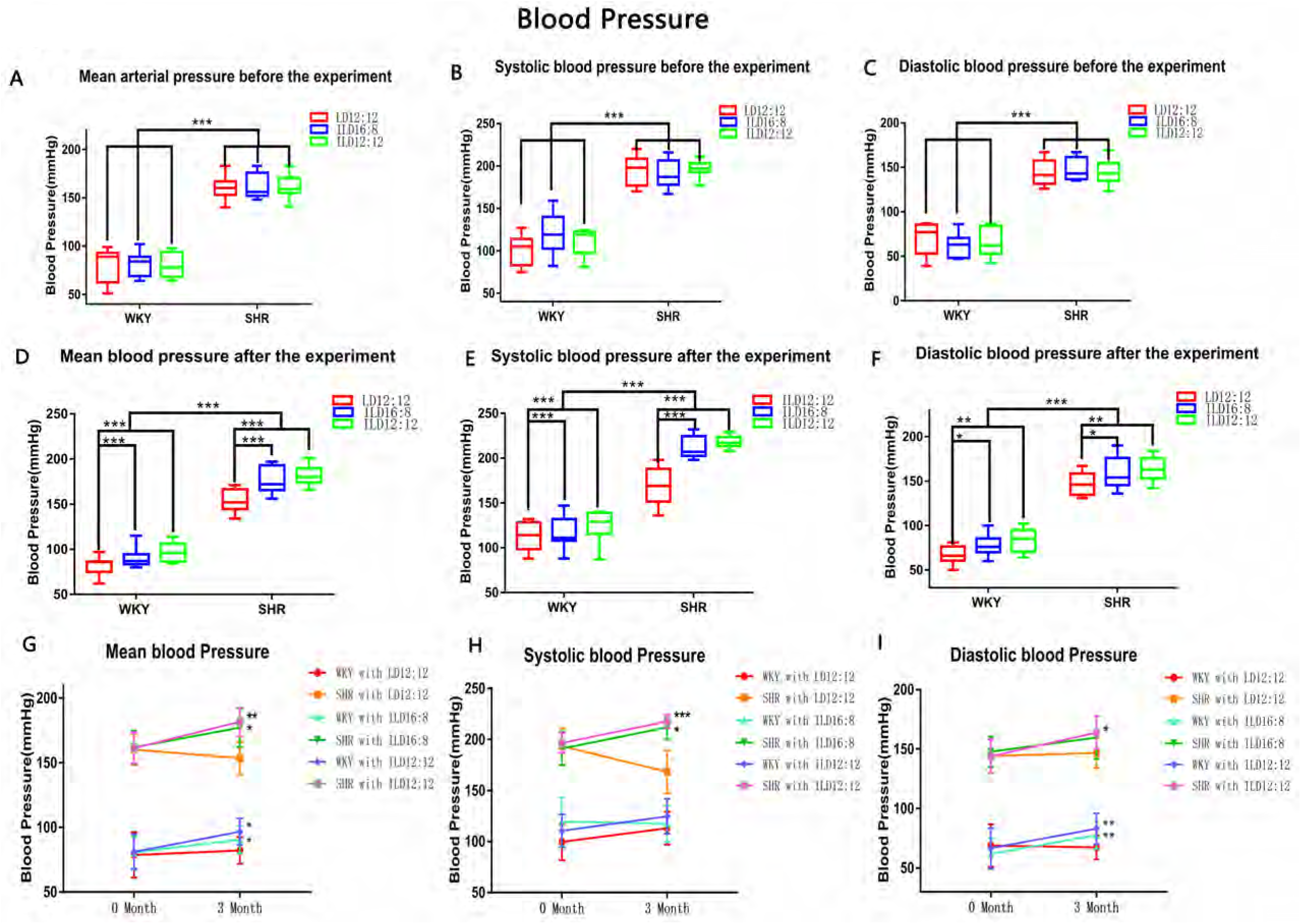
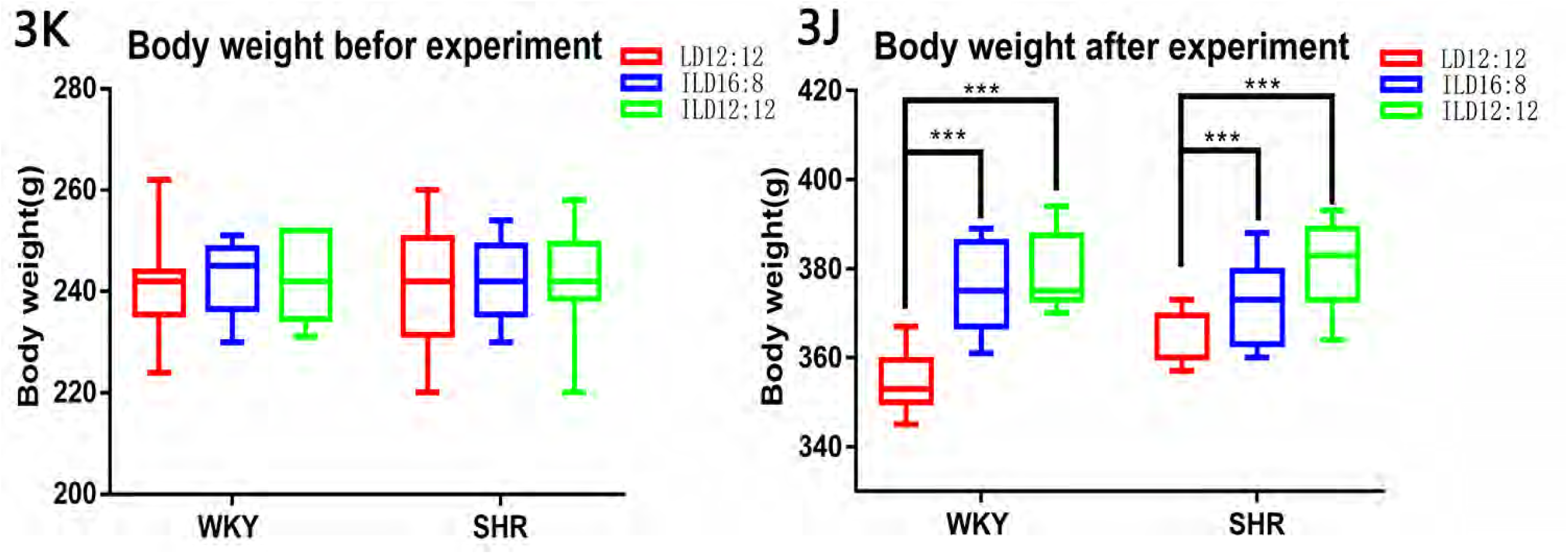
A-C, mean arterial pressure, systolic blood pressure, and diastolic blood pressure data of the six groups of rats before model establishment. D-F, mean arterial pressure, systolic blood pressure, and diastolic blood pressure data of the six groups of rats on day 90 after model establishment. G-I, mean arterial pressure, systolic blood pressure, and diastolic blood pressure data of the six groups of rats before and after model establishment. K body weight data of the six groups of rats before model establishment. J, body weight data of the six groups of rats on day 90 after model establishment. * *p* < 0.05, ** *p* < 0.01, *** *p* < 0.001. A two-factor design was used to compare the six groups of rats. The *post-hoc* Bonferroni was used for comparisons between groups. The paired t-test was used for comparisons of blood pressure before and after model establishment.

Figures 3D–3F represent the statistical results for the mean arterial pressure, systolic blood pressure, and diastolic blood pressure of the rats after the experiment. At 3 months after the establishment of the model, the mean blood pressure (F=16.121, *p*<0.001), systolic blood pressure (F=18.715, *p*<0.001), and diastolic blood pressure (F=7.160, *p*<0.01) in the SW groups were significantly higher than those determined in the LD12:12 groups for both the WKY rats and SHR. There were no significant differences between the ILD16:8 and ILD12:12 groups in mean arterial pressure, systolic blood pressure, and diastolic blood pressure (*p*>0.05). Under the same LD conditions, the mean blood pressure (F=648.406, *p*<0.001), systolic blood pressure (F=357.278, *p*<0.001), and diastolic blood pressure (F=490.554, *p*<0.001) of the SHR were significantly higher than those recorded in the WKY rats.

Figure 3G shows that, at 3 months after the establishment of the model, the mean blood pressure in the SW groups was significantly higher than that measured before the experiment in both the WKY rats and SHR (*p*<0.05). However, the mean blood pressure in SHR(LD12:12) and WKY(LD12:12) rats after the experiment was not significantly different compared with that measured prior to the experiment (*p*>0.05).

Figure 3H shows that, at 3 months after the establishment of the model, the systolic blood pressure in the SHR (ILD16:8) and SHR (ILD12:12) groups was significantly higher compared with that measured prior to the experiment (*p*<0.05). However, the systolic blood pressure in SHR (LD12:12), WKY (LD12:12), WKY (ILD12:12), and WKY (ILD16:8) rats after the experiment was not significantly different versus that recorded before the experiment (*p*>0.05).

Figure 3I shows that, at 3 months after the establishment of the model, the diastolic blood pressure in the SHR (ILD12:12), WKY (ILD16:8), and WKY (ILD12:12) groups was significantly higher compared with that observed before the experiment (*p*<0.05). However, the diastolic blood pressure in SHR (LD12:12), WKY (LD12:12), and SHR (ILD16:8) after the experiment was not significantly different versus that measured prior to the experiment (*p*>0.05).

### 3. Analysis of weight

At 3 months following the establishment of the model, the weight of the SW group was significantly higher than that reported in the LD12:12 group (F=29.716, *p*<0.001) (Figure 3J). There was no significant difference in body weight between the ILD16:8 and ILD12:12 groups (*p*>0.05). Under the same LD conditions, there was no significant difference in the weight between the SHR and WKY rats (F=2.47, *p*>0.05).

### 4. Vascular ultrasound

Internal carotid artery: Figures 4A–E show the intima-media thickness, blood flow velocity, resistance index, systolic diameter, and diastolic diameter of the internal carotid artery. The statistical analysis showed that there was no significant difference in the intima-media thickness (F=0.153, *p*>0.05), blood flow velocity (F=0.473, *p*>0.05), resistance index (F=0.228, *p*>0.05) systolic diameter (F=1.969, *p*>0.05), and diastolic diameter (F=1.472, *p*>0.05) between the SW and LD12:12 groups. However, under the same LD conditions, the blood flow velocity (F=73.025, *p*<0.001) and resistance index (F=4.885, *p*<0.05) in the WKY rats were significantly higher than those observed in the SHR.

**Figure 4:**
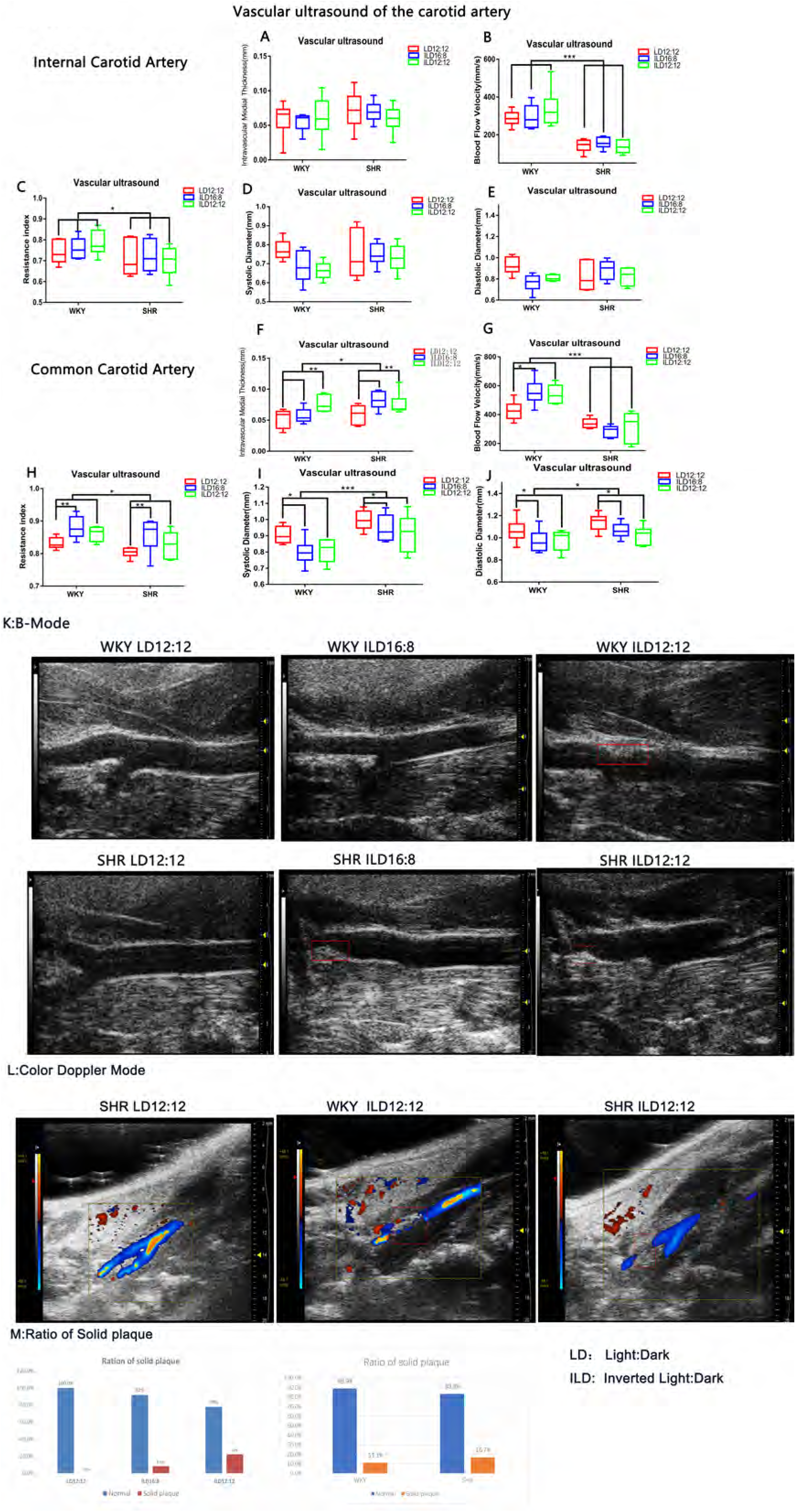
Vascular ultrasound data from the six groups of rats. A-E, intima-media thickness, blood flow velocity, resistance index, systolic diameter, and diastolic diameter of the internal carotid artery, respectively. F-J, intima-media thickness, blood flow velocity, resistance index, systolic diameter, and diastolic diameter of the common carotid artery, respectively. * *p* < 0.05, ** *p* < 0.01, *** *p* < 0.001. A two-factor design was used to compare the six groups of rats. The *post-hoc* Bonferroni was used for comparisons between groups. K, structural diagram of carotid B-mode ultrasound of the six groups of rats. Red squares indicate solid carotid plaques. L, color Doppler imaging of carotid blood flow in the six groups of rats. Red box indicates the area of interrupted carotid blood flow. M, rates of solid plaque formation for the three photoperiods (left) and hypertension factors (right).

Common carotid artery: The intima-media thickness of the ILD12:12 rats was significantly greater than that of the LD12:12 rats (*p* < 0.05) (Figure 4F).However, there was no significant difference in the intima-media thickness of the common carotid artery between the ILD16:8 and ILD12:12 groups (*p*>0.05), or between the LD12:12 and ILD16:8 groups (*p*=0.092). Under the same LD conditions, the common carotid artery intima-media of the SHR was significantly thicker than that observed in the WKY rats (F=4.344, *p*<0.05).

Figure 4G: The two-way ANOVA did not show significant difference in the carotid artery blood flow velocity between the LD12:12 and SW groups (F=1.452, *p*>0.05) (Figure 5G). Under the same LD conditions, the blood flow velocity of the WKY rats was significantly higher than that reported in the SHR (F=67.458, *p*<0.001). However, compared with those in the WKY (LD12:12) rats and SHR (LD12:12), the change in the blood flow velocity in the WKY (SW) rats was different from that in the SHR (SW) rats. Subsequently, one-way ANOVA of the different LDs between the WKY rats and SHR was performed. The results showed that the common carotid artery blood flow velocity in the WKY (ILD16:8) rats was significantly higher than that reported in the WKY (LD12:12) rats (*p*<0.05). However, there was no significant difference in the blood flow velocity of the common carotid artery between the ILD16:8 and ILD12:12 groups (*p*>0.05), or between the LD12:12 and ILD12:12 groups (*p*=0.053). Notably, there was no significant change in the common carotid artery blood flow velocity between the SHR (LD12:12) and SHR (SW) rats (F=0.964, *p*>0.05).

**Figure 5:**
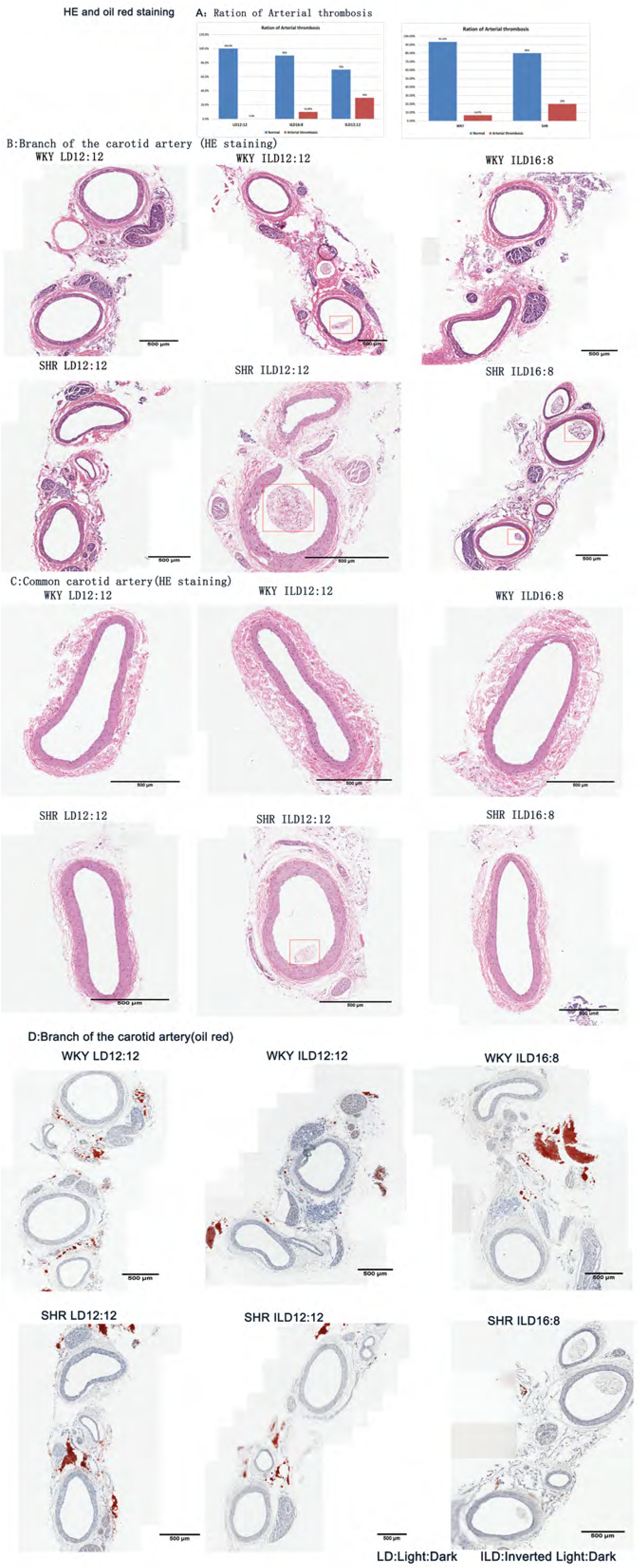
A, rates of arterial thrombosis in the carotid arteries for the three photoperiods (left) and hypertension factors (right). B and C, HE sections of the carotid artery bifurcation (internal, external, and common) and common carotid arteries of the six groups of rats, respectively. Red square indicates carotid arterial thrombosis. Red blood cells, platelets, and fibrin are visible in the arterial thrombus. D, Oil Red O-stained section of the carotid bifurcation in the six groups of rats. No solid atherosclerotic plaques were found on Oil Red O staining of the carotid artery in the six groups of rats.

Figure 4H: Compared with LD12:12, ILD16:8 significantly increased the resistance index of the common carotid arteries (*p*<0.01). There was no significant difference in the resistance index between ILD16:8 and ILD12:12 (*p*>0.05), or between the LD12:12 and ILD12:12 groups (*p*>0.05). However, the resistance index in the WKY rats was significantly higher than that observed in the SHR under the same LD conditions (F=6.439, *p*<0.05).

Figure 4I: Compared with LD12:12, ILD12:12 significantly reduced the systolic diameter of the common carotid artery (*p*<0.05). There was no significant difference in the systolic diameter between ILD16:8 and ILD12:12 (*p*>0.05), or between the LD12:12 and ILD16:8 groups (*p*=0.072). However, the systolic diameter in the SHR was significantly greater than that measured in the WKY rats under the same LD conditions (F=17.963, *p*<0.001).

Figure 4J: Compared with LD12:12, ILD12:12 significantly reduced the diastolic diameter of the common carotid artery (*p*<0.05). There was no significant difference in the diastolic diameter between ILD16:8 and ILD12:12 (*p*>0.05), or between the LD12:12 and ILD16:8 groups (*p*=0.09). However, under the same LD conditions, the diastolic diameter in the SHR rats was significantly greater than that observed in the WKY rats (F=5.74, *p*<0.05).

Figure 4M: Solid plaques were found in the lumen of the carotid artery in some SW groups (Figure 5M). Fisher’s exact test did not show significant difference in the rate of solid plaque formation between the SW and LD12:12 groups (*p*>0.05). However, the rate of solid plaque formation in the ILD12:12 group was higher than that recorded in the LD12:12 group. The solid plaques in the carotid artery resulted in partial interruption of blood flow (Figure 5L). Examination using vascular ultrasound was unable to determine the nature of these solid plaques; this must be investigated via carotid artery pathology examination.

### 5. Carotid arterial sections

HE staining: HE sections showed that some of the solid plaques found in the carotid artery of the SW groups contained intact or ruptured red blood cells, platelets, and fibrin. Therefore, these solid plaques were arterial thrombi (Figure 5B). Although Figure 5A shows that there was no significant difference in the rate of formation of arterial thrombosis between the SW and LD12:12 groups (*p*>0.05), the mean rate of arterial thrombosis in the SW group was higher than that reported in the LD12:12 group. Under the same LD conditions, the rate of formation of arterial thrombosis in the SHR was not significantly different from that determined in the WKY rats (*p*>0.05).

Figure 5D: Oil Red O staining: Fat droplets or atherosclerotic plaques were not found in the walls of the carotid arteries of the rats.

Masson’s trichrome staining: Figure 6A shows that the SW group had a significantly increased intima-media thickness of the internal carotid artery compared with the LD12:12 group (F=12.252, *p*<0.001). The difference in the intima-media thickness between the ILD16:8 and ILD12:12 groups was not significant (*p*>0.05). The intima-media thickness of the internal carotid artery in the SHR was significantly greater than that measured in the WKY rats under the same LD conditions (F=10.892, *p*<0.01). Figure 6B shows that the SW group had a significantly increased intima-media thickness of the common carotid artery compared with the LD12:12 group (F=20.39, *p*<0.001). The difference in the intima-media thickness between the ILD16:8 and ILD12:12 groups was not significant (*p*>0.05). The intima-media thickness of the common carotid artery in the SHR was significantly greater than that reported in the WKY rats under the same LD conditions (F=38.537, *p*<0.001).

**Figure 6:**
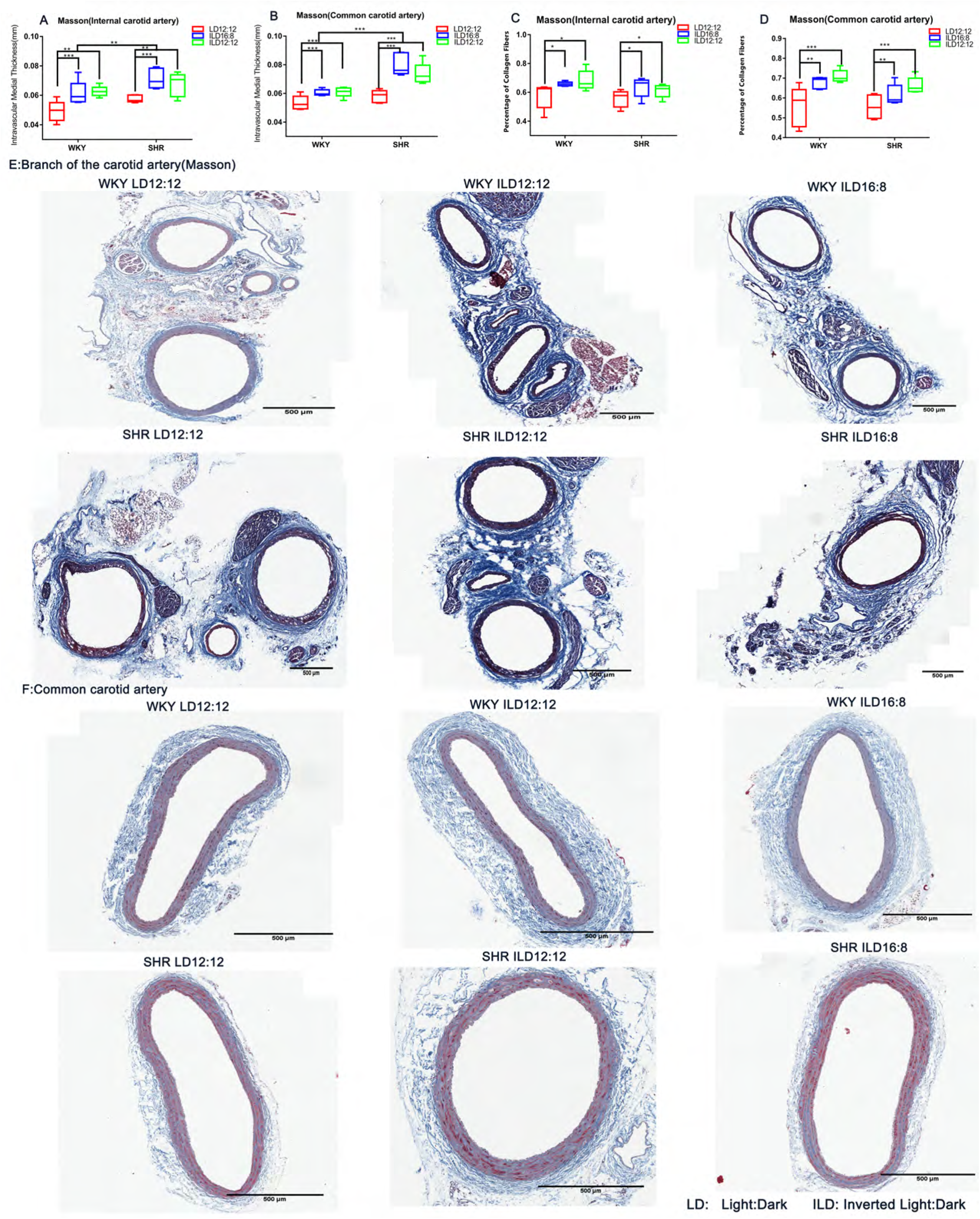
A and B, internal and common carotid artery intima-media thickness data of the six groups of rats, respectively. C and D, internal and common carotid artery collagen fiber ratios of the six groups of rats, respectively. * *p* < 0.05, ** *p* < 0.01, *** *p* < 0.001. A two-factor design was used to compare the six groups of rats. The *post-hoc* Bonferroni was used for comparisons between groups. E and F, Masson-stained sections of the internal carotid artery and common carotid artery in the six groups of rats, respectively.

Figure 6C shows that the SW group had a significantly increased collagen fiber ratio in the internal carotid artery compared with the LD12:12 group (F=5.692, *p*<0.01). There was no significant difference in the collagen fiber ratio between the ILD16:8 and ILD12:12 groups (*p*>0.05). Under the same LD conditions, there was no significant difference in the carotid artery collagen fiber ratio between the two groups (F=2.711, *p*>0.05). Figure 6D shows that the SW group had a significantly increased collagen fiber ratio in the common carotid artery compared with the LD12:12 group (F=13.501, *p*<0.001). There was no significant difference in the collagen fiber ratio between the ILD16:8 and ILD12:12 groups (*p*>0.05). There was also no significant difference in the internal carotid artery collagen fiber ratio between the SHR and WKY rats under the same LD conditions (F=3.092, *p*>0.05).

AGTR: Figure 7A shows that the SW group had a significantly increased total AGTR density in the common carotid artery compared with the LD12:12 group (F=4.778, *p*<0.05). The difference in the total AGTR density in the common carotid artery between the ILD16:8 and ILD12:12 groups was not significant (*p*>0.05). The total AGTR density in the common carotid artery of the SHR was not significantly different from that measured in the WKY rats under the same LD conditions (F=1.216, *p*>0.05).

**Figure 7:**
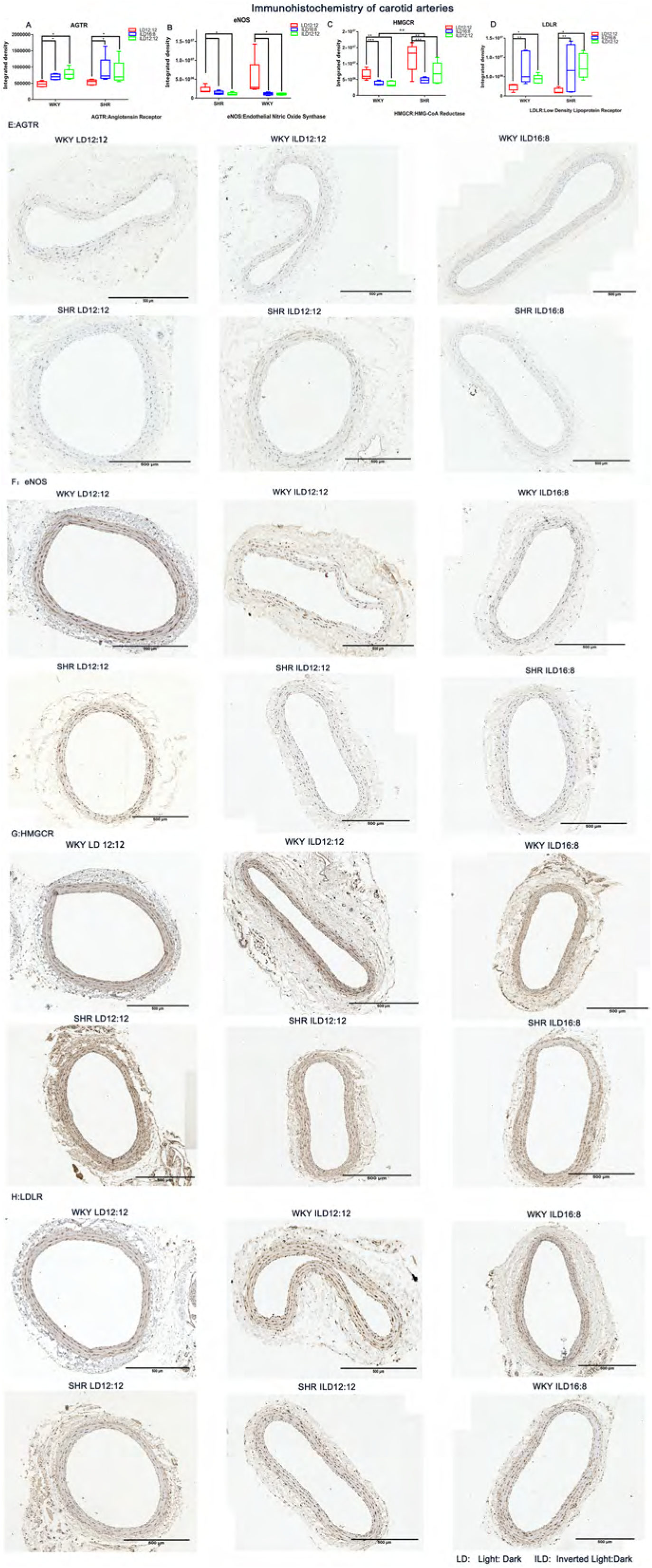
A-D, AGTR, eNOS, HMGCR, and LDLR immunohistochemistry data for the common carotid artery in the six groups of rats, respectively. * *p* < 0.05, ** *p* < 0.01, *** *p* < 0.001. A two-factor design was used to compare the six groups of rats. The *post-hoc* Bonferroni was used for comparisons between groups. E-H, AGTR, eNOS, HMGCR, and LDLR immunohistochemical sections of the common carotid artery in the six groups of rats, respectively.

eNOS: Figure 7B shows that the ILD12:12 group had a significantly reduced total eNOS density in the common carotid artery compared with the LD12:12 group (*p*<0.05). However, there was no difference in the total eNOS density in the common carotid artery between the ILD16:8 and ILD12:12 groups (*p*>0.05), or between the LD12:12 and ILD16:8 groups (*p*=0.068). The total eNOS density in the common carotid artery of the SHR was not significantly different from that recorded in the WKY rats under the same LD conditions (F=1.194, *p*>0.05).

HMGCR: Figure 7C shows that the SW group had a significantly reduced total HMGCR density in the common carotid artery compared with the LD12:12 group (F=10.36, *p*<0.001). The difference in the total HMGCR density in the common carotid artery between the ILD16:8 and ILD12:12 groups was not significant (*p*>0.05). However, the total HMGCR density in the common carotid artery of the SHR was higher than that observed in the WKY rats under the same LD conditions (F=10.335, *p*<0.01).

LDLR: Figure 7D shows that the SW group had a significantly increased total LDLR density in the common carotid artery compared with the LD12:12 group (F=6.887, *p*<0.01). The difference in the total LDLR density in the common carotid artery between the ILD16:8 and ILD12:12 groups was not significant (*p*>0.05). The total LDLR density in the common carotid artery of the SHR was not significantly different from that reported in the WKY rats under the same LD conditions (F=0.372, *p*>0.05).

Gross Oil Red O staining: Gross Oil Red O staining showed that some SW groups developed atherosclerotic plaques in the abdominal aorta, which appeared orange-red (Figure 8B). The number and area of atherosclerotic plaque formation in the SW groups are shown in Table 1C. The number and area of atherosclerotic plaque formation in the SW groups are shown in Table 1C. Figure 8C showed that the area ratio of positive plaque in different photoperiods was not exactly the same (F = 3.774, *p* < 0.05). However, the difference in the area ratio of positive plaque between the ILD12:12 and ILD16:8 groups was not significant (*p* = 0.086). The difference in the area ratio of positive plaque between the LD12:12 and ILD12:12 groups was not significant (*p* = 0.086). The difference in the area ratio of positive plaque between the LD12:12 and ILD16:8 groups was not significant (p = 1). Under the same LD conditions, there was no significant difference in the rate of atherosclerotic plaque formation between the SHR and WKY rats (F=1.302, *p*>0.05).

**Figure 8:**
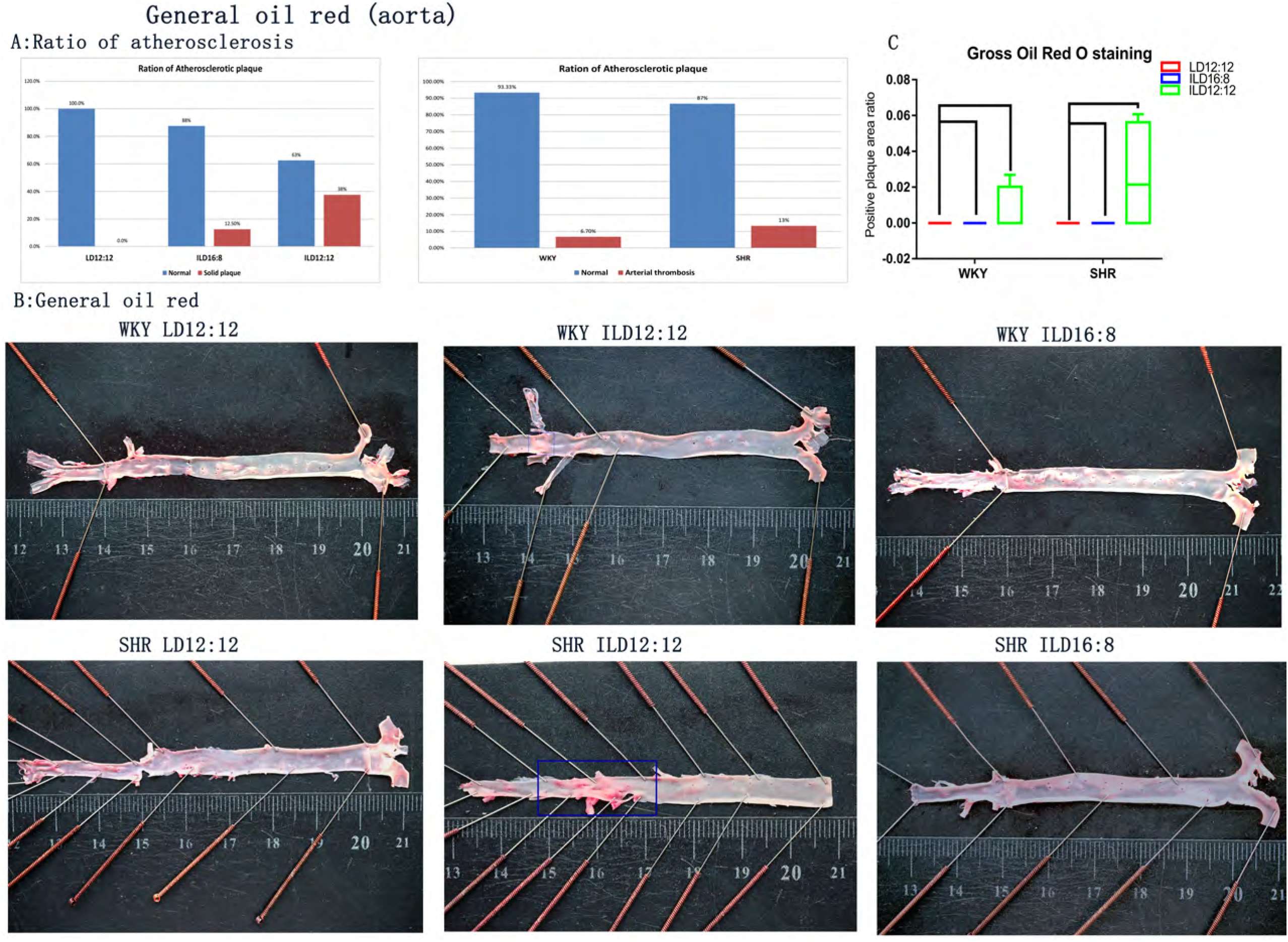
A, rates of aortic atherosclerotic plaque formation for the three LDs (left) and hypertension factors (right). B, gross Oil Red O staining of a longitudinal section of the aorta in the six groups of rats. Red squares indicate atherosclerotic plaque formation. C, the area ratio of positive plaque data of the six groups of rats. A two-factor design was used to compare the six groups of rats. The *post-hoc* Bonferroni was used for comparisons between groups.

**Table 1:**
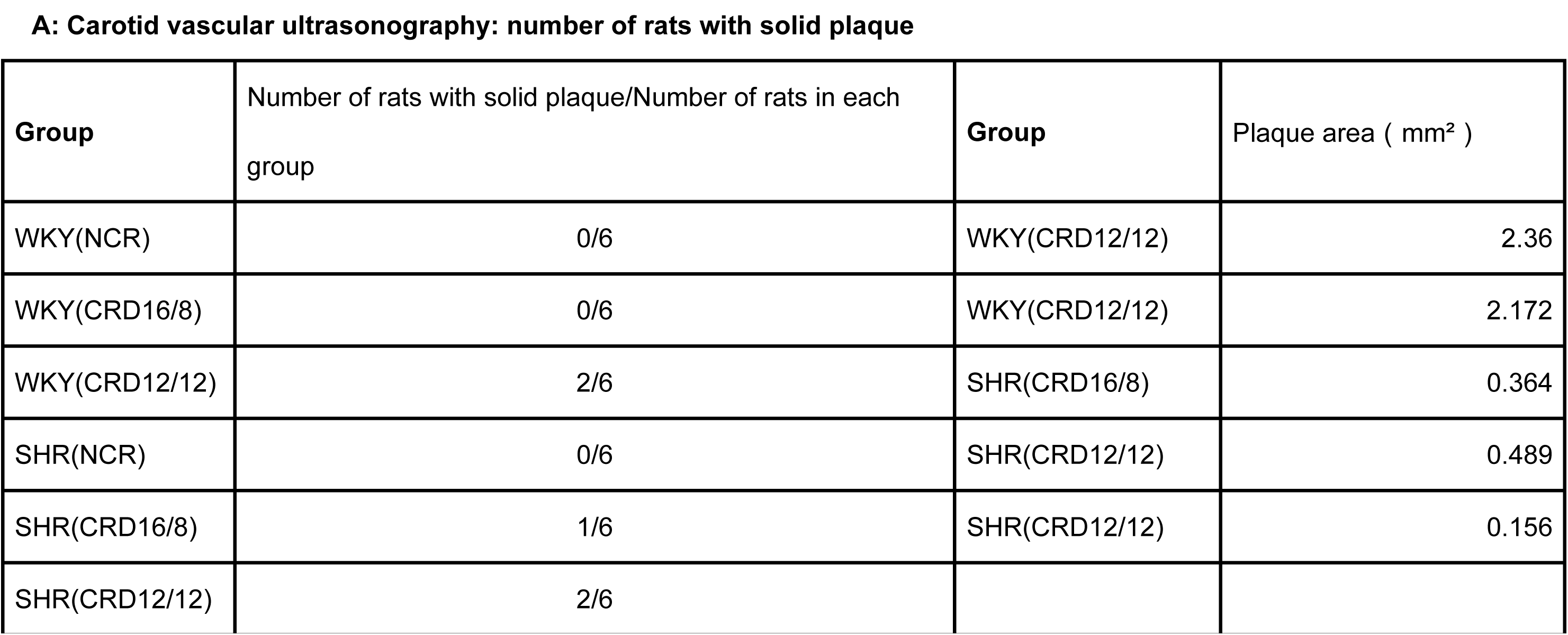
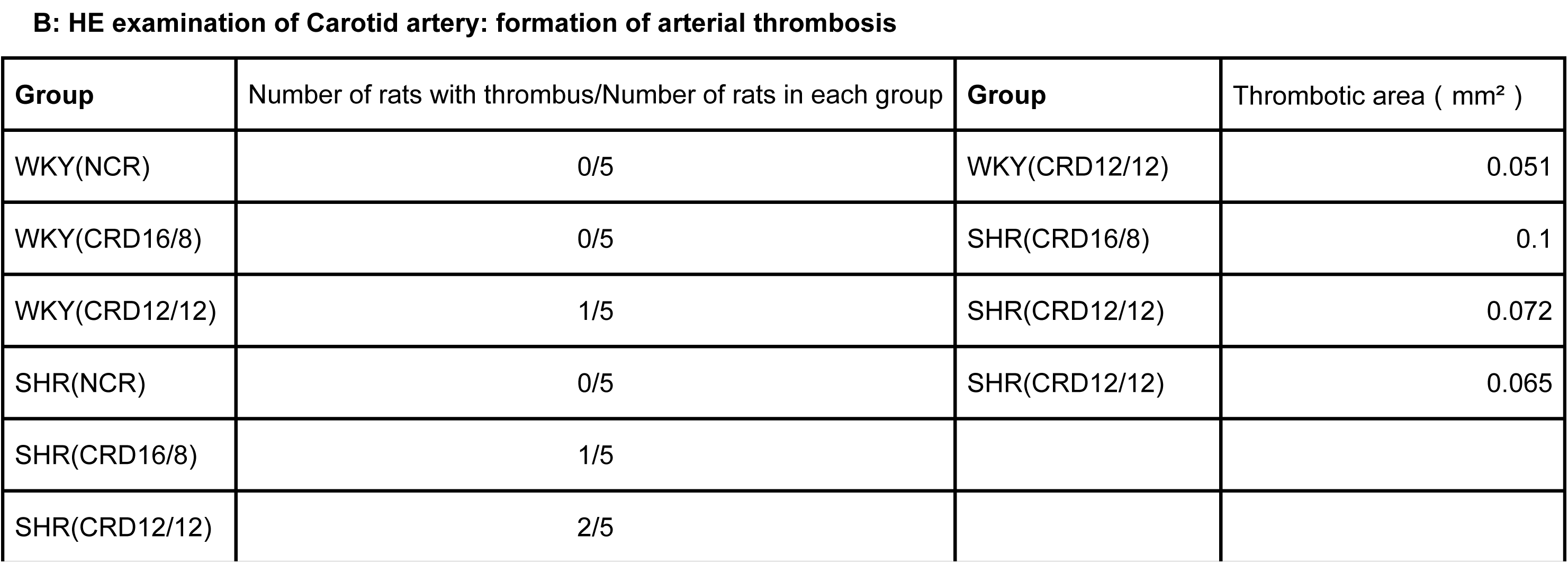
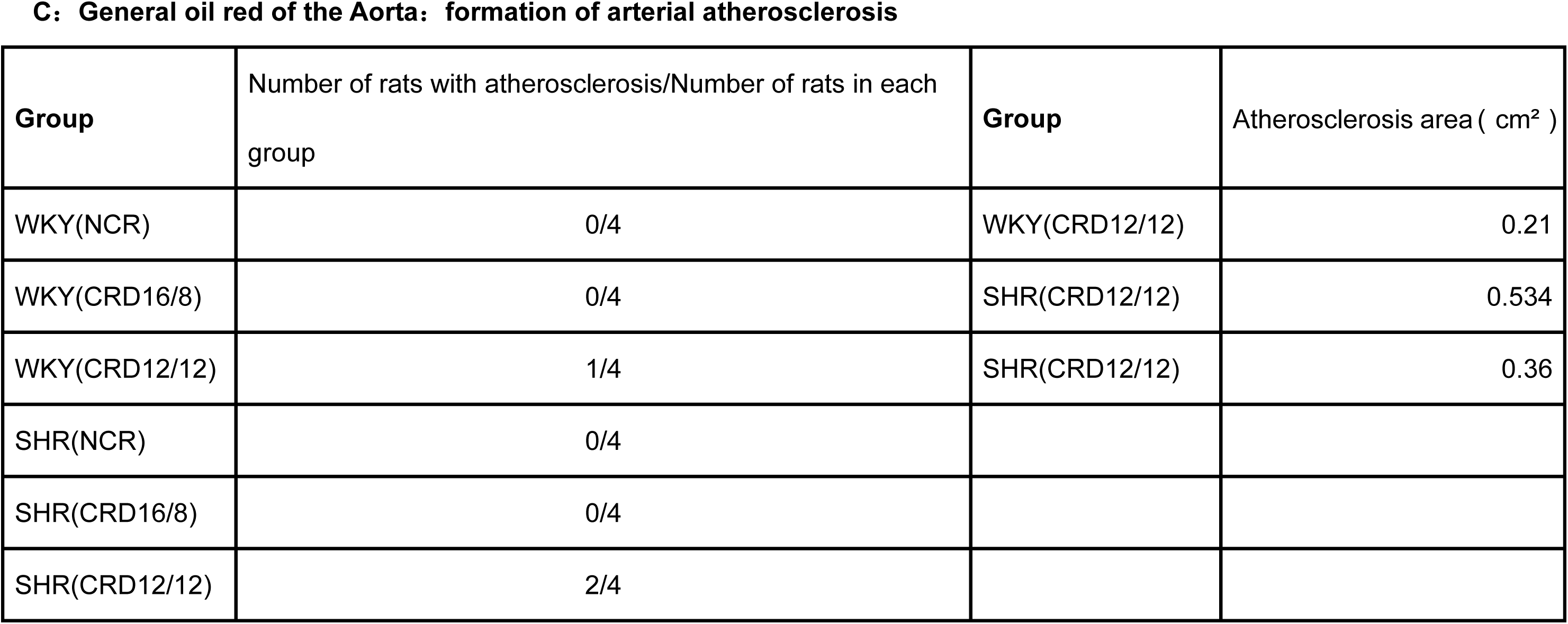
A, rates of solid plaque formation and area of solid plaques in the carotid arteries of the six groups of rats. B, rates of carotid arterial thrombosis and the area of arterial thrombosis in the six groups of rats. C, rates of aortic atherosclerotic plaque formation and the area of atherosclerotic plaques in the six groups of rats.

| **Group** | Number of rats with solid plaque/Number of rats in each group | **Group** | Plaque area ( mm<sup>2</sup> ) |
| --- | --- | --- | --- |
| WKY(NCR) | 0/6 | WKY(CRD12/12) | 2.36 |
| WKY(CRD16/8) | 0/6 | WKY(CRD12/12) | 2.172 |
| WKY(CRD12/12) | 2/6 | SHR(CRD16/8) | 0.364 |
| SHR(NCR) | 0/6 | SHR(CRD12/12) | 0.489 |
| SHR(CRD16/8) | 1/6 | SHR(CRD12/12) | 0.156 |
| SHR(CRD12/12) | 2/6 |  |  |

**B: HE examination of Carotid artery: formation of arterial thrombosis**
| **Group** | Number of rats with thrombus/Number of rats in each group | **Group** | Thrombotic area ( mm<sup>2</sup> ) |
| --- | --- | --- | --- |
| WKY(NCR) | 0/5 | WKY(CRD12/12) | 0.051 |
| WKY(CRD16/8) | 0/5 | SHR(CRD16/8) | 0.1 |
| WKY(CRD12/12) | 1/5 | SHR(CRD12/12) | 0.072 |
| SHR(NCR) | 0/5 | SHR(CRD12/12) | 0.065 |
| SHR(CRD16/8) | 1/5 |  |  |
| SHR(CRD12/12) | 2/5 |  |  |

**C: General oil red of the Aorta: formation of arterial atherosclerosis**
| **Group** | Number of rats with atherosclerosis/Number of rats in each group | **Group** | Atherosclerosis area ( cm<sup>2</sup> ) |
| --- | --- | --- | --- |
| WKY(NCR) | 0/4 | WKY(CRD12/12) | 0.21 |
| WKY(CRD16/8) | 0/4 | SHR(CRD12/12) | 0.534 |
| WKY(CRD12/12) | 1/4 | SHR(CRD12/12) | 0.36 |
| SHR(NCR) | 0/4 |  |  |
| SHR(CRD16/8) | 0/4 |  |  |
| SHR(CRD12/12) | 2/4 |  |  |

### 6. Serology

IL-6: The levels of IL-6 in the serum of the ILD12:12 group were significantly increased compared with those measured in the LD12:12 group (*p*<0.05) (Figure 9A) and in the ILD16:8(p<0.05). However, there was no significant difference in the levels of IL-6 between the LD12:12 and ILD16:8 (*p*>0.05). Under the same LD conditions, there was no significant difference in the levels of IL-6 between the SHR and WKY rats (F=1.517, *p*>0.05).

**Figure 9:**
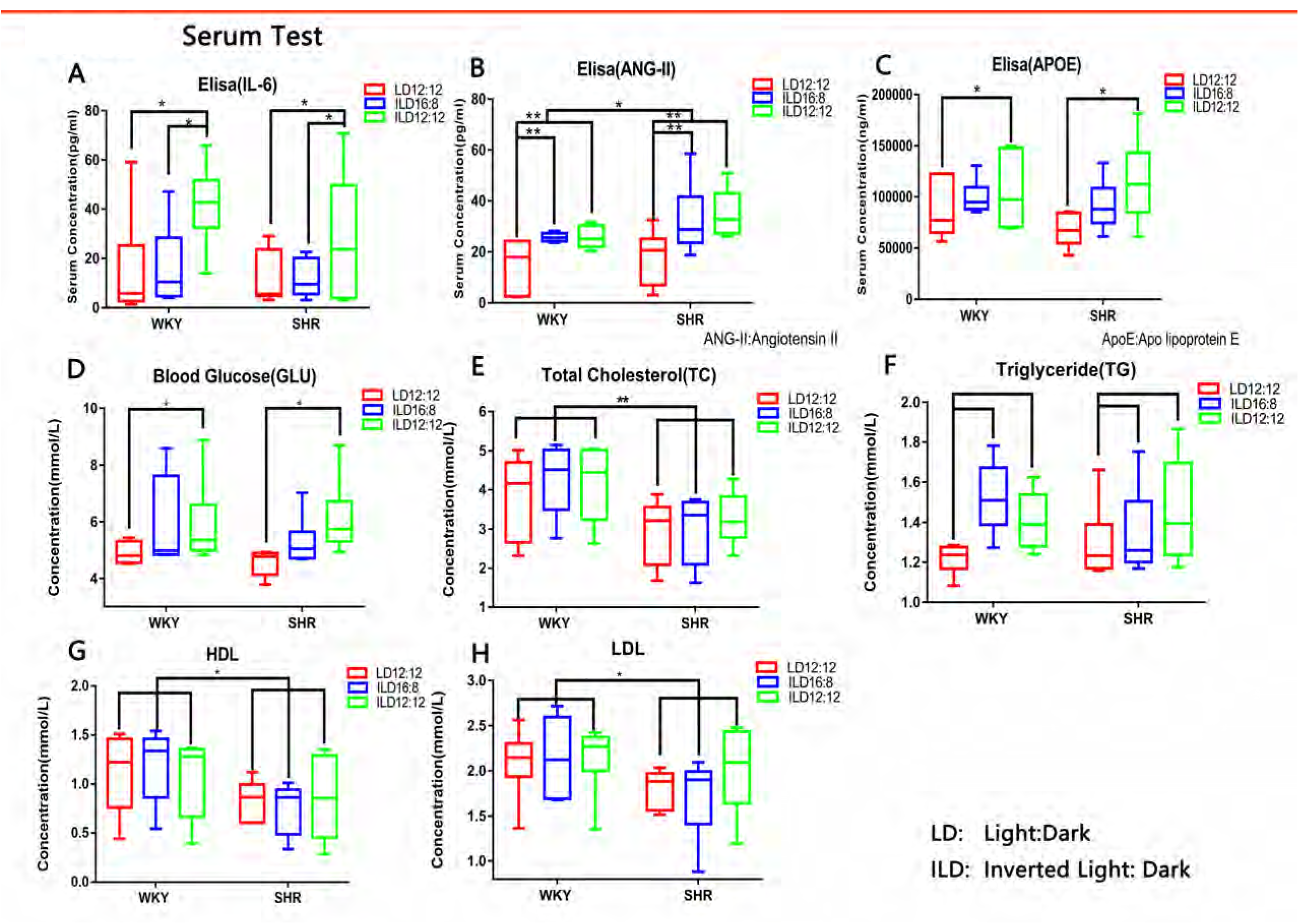
A-H, serum IL-6, ANG-II, APOE, Glu, TC, TG, HDL, and LDL data of the six groups of rats. * *p* < 0.05, ** *p* < 0.01, *** *p* < 0.001. A two-factor design was used to compare the six groups of rats. The *post-hoc* Bonferroni was used for comparisons between groups.

ANG-II: The levels of ANG-II in the SW group were significantly increased compared with those recorded in the LD12:12 group (F=8.362, *p*<0.01) (Figure 9B). However, there was no significant difference in the levels of ANG-II between the ILD16:8 and ILD12:12 groups (*p*>0.05). Under the same LD conditions, the levels of ANG-II in the SHR were significantly higher than those reported in the WKY rats (F=4.287, *p*<0.05).

APOE: The levels of APOE in the serum of the ILD12:12 group were significantly increased compared with those observed in the LD12:12 group (*p*<0.05) (Figure 9C). However, there was no significant difference in the levels of APOE between the ILD16:8 and ILD12:12 groups (*p*>0.05), or between the LD12:12 and ILD16:8 groups (*p*>0.05). Under the same LD conditions, there was no significant difference in the levels of APOE between the SHR and WKY rats (F=0.366, *p*>0.05).

Blood Glu: The levels of blood Glu in the ILD12:12 group were significantly increased compared with those recorded in the LD12:12 group (*p*<0.05) (Figure 9D). However, there was no significant difference in the levels of blood Glu between the ILD16:8 and ILD12:12 groups (*p*>0.05), or between the LD12:12 and ILD16:8 groups (*p*>0.05). Under the same LD conditions, there was no significant difference in the levels of blood Glu between the SHR and WKY rats (F=0.428, *p*>0.05).

TG: The levels of TG in different photoperiods were not exactly the same (F=3.564, *p*<0.05) (Figure 9F). However, the difference in the levels of TG between the ILD12:12 and ILD16:8 groups was not significant (*p* >0.05). There was no significant difference in the levels of TG between the ILD12:12 and LD12:12 groups (*p*=0.087), or between the LD12:12 and ILD16:8 groups (*p*=0.08). Under the same LD conditions, there was no significant difference in the levels of TG between the SHR and WKY rats (F=0.086, *p*>0.05).

Figures 9E, 9G, and 9H show the levels of TC, HDL, and LDL, respectively, in the serum. There were no significant differences in TC (F=0.434, *p*>0.05), HDL (F=0.005, *p*>0.05), and LDL (F=0.473, *p*>0.05) between the SW and LD12:12 groups. However, under the same LD conditions, the levels of TC (F=11.832, *p*<0.01), HDL (F=7.02, *p*<0.05), and LDL (F=4.476, *p*<0.05) were lower in the SHR than in the WKY rats.

These experiments showed that the two types of SW significantly increased the levels of ANG-II in the serum. ILD12:12 markedly increased the levels of IL-6, GLU, and APOE. ILD16:8 resulted in an increasing trend for the levels of IL-6 and APOE; however, the difference was not significant. In addition, the two types of SW did not significantly affect the levels of TC, HDL, and LDL.

### 7. ASL examination

Figure 10A shows that the SW group had a significantly reduced CBF compared with the LD12:12 group (F=8.426, *p*<0.01). There was no significant difference in blood flow between the ILD16:8 and ILD12:12 groups (*p*>0.05). Moreover, there was no significant difference in the hippocampal blood flow between the SHR and WKY rats under the same LD conditions (F=0.454, *p*>0.05).

**Figure 10:**
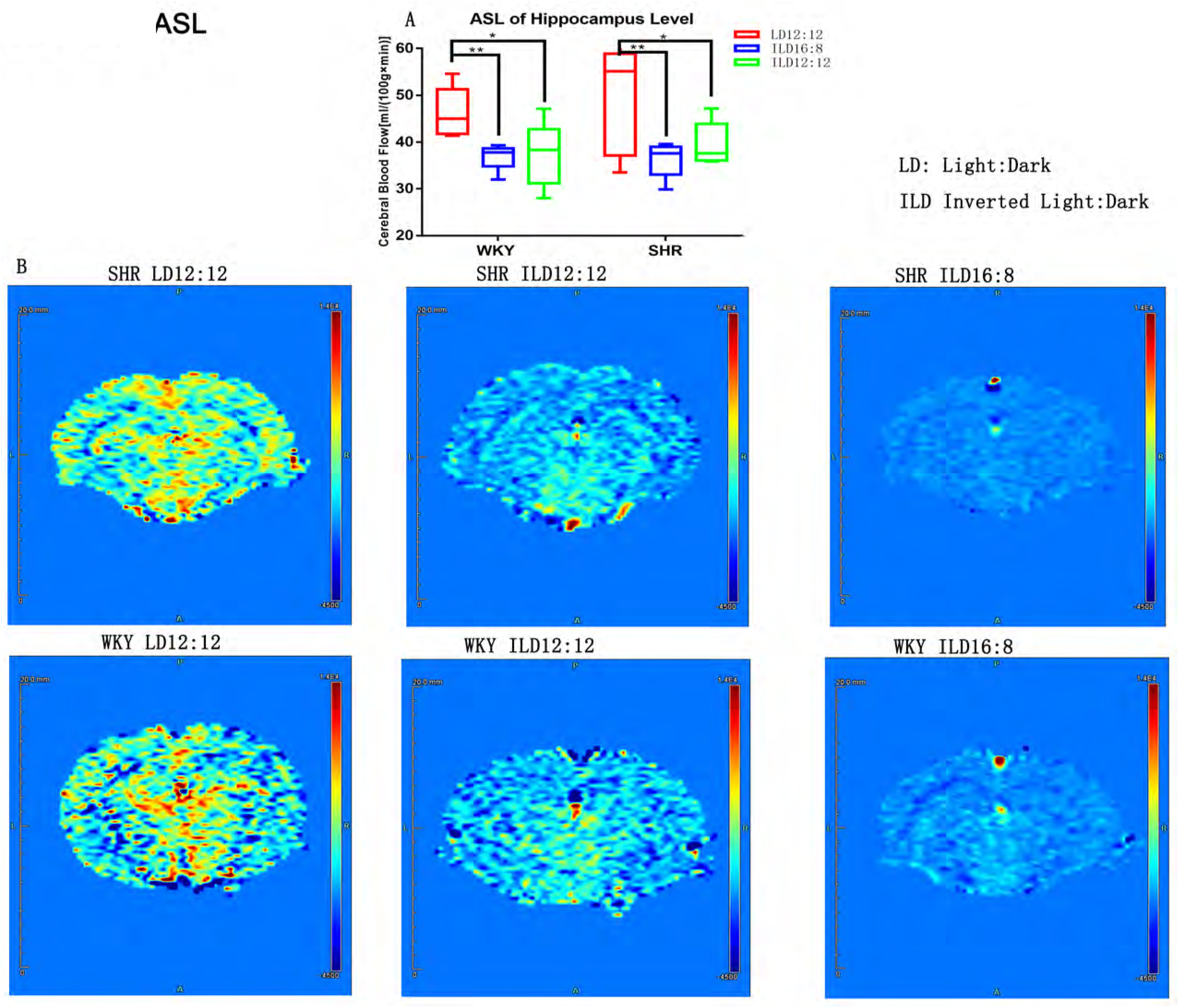
A, CBF value data on hippocampal plane ASL examination for the six groups of rats. * *p* < 0.05, ** *p* < 0.01, *** *p* < 0.001. A two-factor design was used to compare the six groups of rats. The *post-hoc* Bonferroni was used for comparisons between groups. B, CBF images of the six groups of rats. Red and yellow areas indicate high CBF, and blue area represents low CBF. CBF, cerebral blood flow.

## Discussion

This study found that both the WKY and the SHR LD12:12 groups did not have vascular lesions. However, a number of rats in the SW groups exhibited lesions in the carotid arteries or aorta. These lesions included carotid intima-media thickening, atherosclerotic plaque formation in the abdominal aorta, and carotid arterial thrombosis. Of note, these changes decreased CBF. These findings suggest that SW is directly associated with arterial diseases in rats. Although there was no formation of atherosclerotic plaque, as shown by Oil Red O staining of the carotid artery in this study, we believe that SW may induce atherosclerotic lesions on the carotid arteries based on the following two observations. Firstly, we detected intima-media thickening of the carotid artery through vascular ultrasound examination and HE staining of tissue sections. Numerous studies have shown that intima-media thickening of the carotid artery is a prodromal lesion of carotid arterial atherosclerosis[^33–36^]. Secondly, we found that the SW groups developed obvious atherosclerotic plaques in the abdominal aorta (Figure 10B). According to the literature, atherosclerosis develops earlier in the abdominal aorta than in other segments of the aorta [^37^]. The biochemistry of carotid arterial blood is similar to that of the aorta, and the two arteries are anatomically similar. Hence, the formation of atherosclerotic plaques in the aorta indicates that such plaques may also form in the carotid artery. Moreover, carotid artery atherosclerotic plaques could not be observed due to insufficient observation time. Based on the results of this study, the development of atherosclerosis may be primarily attributed to the following two points. Firstly, SW-induced metabolic disorders may damage vascular endothelial cells via multiple pathways. Increased levels of ANG-II in the serum and expression of AGTR in the vascular wall, caused by SW-triggered intima-media thickening and increased collagen fibers in the vascular wall [^38–43^], ultimately resulted in elevated blood pressure. Combined with hyperglycemia, hyperlipidemia, and high levels of IL-6 caused by SW, elevated blood pressure may cause significant damage to vascular endothelial cells [^43^]. Secondly, decreased expression of HMGCR in the vascular wall and increased expression of LDLR may lead to abnormal deposition of lipids in the vascular wall [^44–46^].

The present study found that the SW groups developed disorders of lipid metabolism and Glu metabolism as follows: (1) the rats gained weight; and (2) the levels of blood Glu, serum TG, APOE, and vascular-wall LDLR were increased, whereas the levels of HMGCR were decreased. LD12:12 maintains the physiological function of the circulatory system[^47, 48^], ensuring the presence of normal and organized structures and normal function of the circulatory system. Under a SW condition, the following adverse changes occur in the body: (1) the nutrient metabolism of the body is no longer synchronized with the activity rhythm; and (2) the nutrient metabolism of the body is no longer synchronized with endocrine hormone secretion [^49^], which may also trigger a variety of disorders related to hormone secretion[^49.50^]. Numerous studies confirmed that disorders of lipid metabolism and Glu metabolism are independent risk factors of atherosclerosis [^51–55^]. Therefore, we suggest that the intima-media thickening of the carotid artery and the atherosclerotic plaques in the abdominal aorta observed in the SW groups of the present study may be associated with disorders of lipid metabolism and Glu metabolism. Firstly, the principal adverse effect of elevated levels of blood Glu and dyslipidemia is the loss of homeostasis [^56^]. In this study, changes in proteins associated with lipid metabolism, including the downregulation of HMGCR expression and upregulation of LDLR expression in the vessel wall, became important “trigger points” for changes in vascular structure and function[^57–59^]. Secondly, SW increased the blood pressure in the WKY rats and exacerbated the blood pressure in the SHR. This is mainly attributed to the increase in the levels of ANG-II in the serum caused by metabolic disorders, as well as increased expression of AGTR and decreased expression of eNOS in the carotid artery, ultimately resulting in increased blood pressure[^60–62^]. In addition, metabolic disorders trigger an elevation in the levels of IL-6, resulting in an increase in the inflammatory response of the blood vessel [^63, 64^]. An article published in Nature in 2006 clearly proposed the concept of metabolic inflammation, which is essentially the enhancement of the body’s inflammatory response caused by metabolic disorders [^65^]. In addition to the effects of the metabolic disorders, metabolic inflammation renders SW even more harmful to the vessel wall. Surprisingly, with regard to metabolic disorders, we observed elevated levels of TG in the SW groups without significant changes to the levels of cholesterol. However, based on the following two points, we suggest that SW may affect the metabolism of cholesterol. Firstly, rats lack a gallbladder [^66^]. Thus, they belong to a group of animals with specific disorders of cholesterol synthesis. Secondly, the immunohistochemical results of the carotid artery showed that SW reduced the expression of HMGCR, indicating that SW can induce disorders of cholesterol metabolism in the vascular wall. Therefore, whether SW elevates the levels of cholesterol should be investigated in future clinical trials.

Vascular ultrasound of the carotid artery revealed that the diameter of the common carotid artery was reduced, and the resistance index increased in both WKY rats and SHR under SW conditions. We believe that these observations are primarily attributed to the following. Firstly, increased intima-media thickness of the carotid artery narrows the vessel, thereby reducing the area for blood flow. Secondly, SW can reduce the compliance of the carotid artery via metabolic disorders. In the present study, the decrease in compliance was primarily associated with the increase in the content of collagen fiber in the vessel wall[^67–69^], and the imbalance in the ratio of eNOS to AGTR expression in the vessel wall. [^60–62^] However, with respect to the blood flow velocity, the effects of SW on the carotid arteries of the two types of rats were not identical. In the WKY rats, SW reduced the diameter of the carotid artery and increased the blood flow velocity. Although SW also reduced the diameter of the carotid artery in the SHR, the diameter in this type of rats was significantly larger than that observed in the WKY rats (Figure 5G). Moreover, the blood flow velocity did not change significantly. The increased blood flow velocity in the WKY rats appeared to be a compensatory response to the narrowing of the vessel diameter, with the main goal of ensuring normal CBF. In the SHR, the diameter of the vessel and blood flow velocity did not change significantly. This is clearly a form of decompensation, as the decline was more obvious from the perspective of reduced blood supply from the carotid artery. In addition, pathological changes in blood flow velocity in the SHR (SW) rats are more likely to cause ischemic stroke [^70, 71^]. This demonstrates that, in the context of SW, hypertension exerts an exacerbating effect on rats.

A very important finding in the present study is that carotid arterial thrombosis was found in some SW groups. However, the statistical results do not currently support the notion that SW can significantly increase the probability of arterial thrombosis. Notably, rats in the LD12:12 group did not have thrombosis, and the rats that formed thrombi were all subjected to SW. Therefore, we highly suspect that SW induced the formation of arterial thrombosis. Expansion of the sample and extension of the modeling time are required to continue observation. According to the current literature, the cause of arterial thrombosis may be related to the following factors. Firstly, examination through vascular ultrasound showed changes in the hemodynamics of the carotid arteries in the SW groups. These presented as altered hemodynamics caused by SW, rapidly increased blood flow velocity in the WKY rats, and decreased vessel diameter and increased resistance index of the carotid arteries in both the WKY rats and SHR [^72–75^]. Secondly, serological examination showed that SW may elevate the levels of IL-6, blood Glu, and ANG-II, and reduce the levels of eNOS. These effects lead to damages in the endothelial cells of the carotid artery and cause the formation of arterial thrombosis [^76–84^]. Thirdly, increased blood pressure induced by SW may also cause the formation of arterial thrombosis [^85, 86^]. Therefore, the development of carotid arterial thrombosis is a result of the widespread adverse effects of SW.

In our subsequent analysis, we found that the adverse effects of SW on the carotid artery are not limited to the carotid artery itself. The carotid artery plays an indispensable role in cerebral blood supply. Therefore, pathological changes in the carotid artery will ultimately affect CBF. Figure 11 shows multiple pathological events that reduce blood supply to the brain via the carotid artery. Firstly, intima-media thickening of the carotid artery may lead to stenosis of the vascular lumen. Secondly, arterial thrombosis may lead to partial interruption of blood supply via the carotid artery. Thirdly, hemodynamic changes may directly alter CBF. Based on these points, we propose that SW interferes with CBF in rats through anatomical and physiological effects on the carotid artery [^87, 88^]. The ASL sequence is a non-invasive, simple, and safe technique to examine CBF [^89^]. Hence, we used this method to measure CBF in the rats. The observed decrease in CBF also confirmed that SW may cause carotid arterial pathology, which consequently leads to decompensation of carotid arterial structure and function. This indicates that the negative changes in carotid arterial structure are associated with the loss of regulatory function and decrease in CBF caused by SW. In contrast to carotid arterial pathologies, SW is more life-threatening owing to the insufficient blood supply to the brain caused by carotid arterial lesions. Therefore, we plan to further investigate this phenomenon in the future.

**Figure 11:**
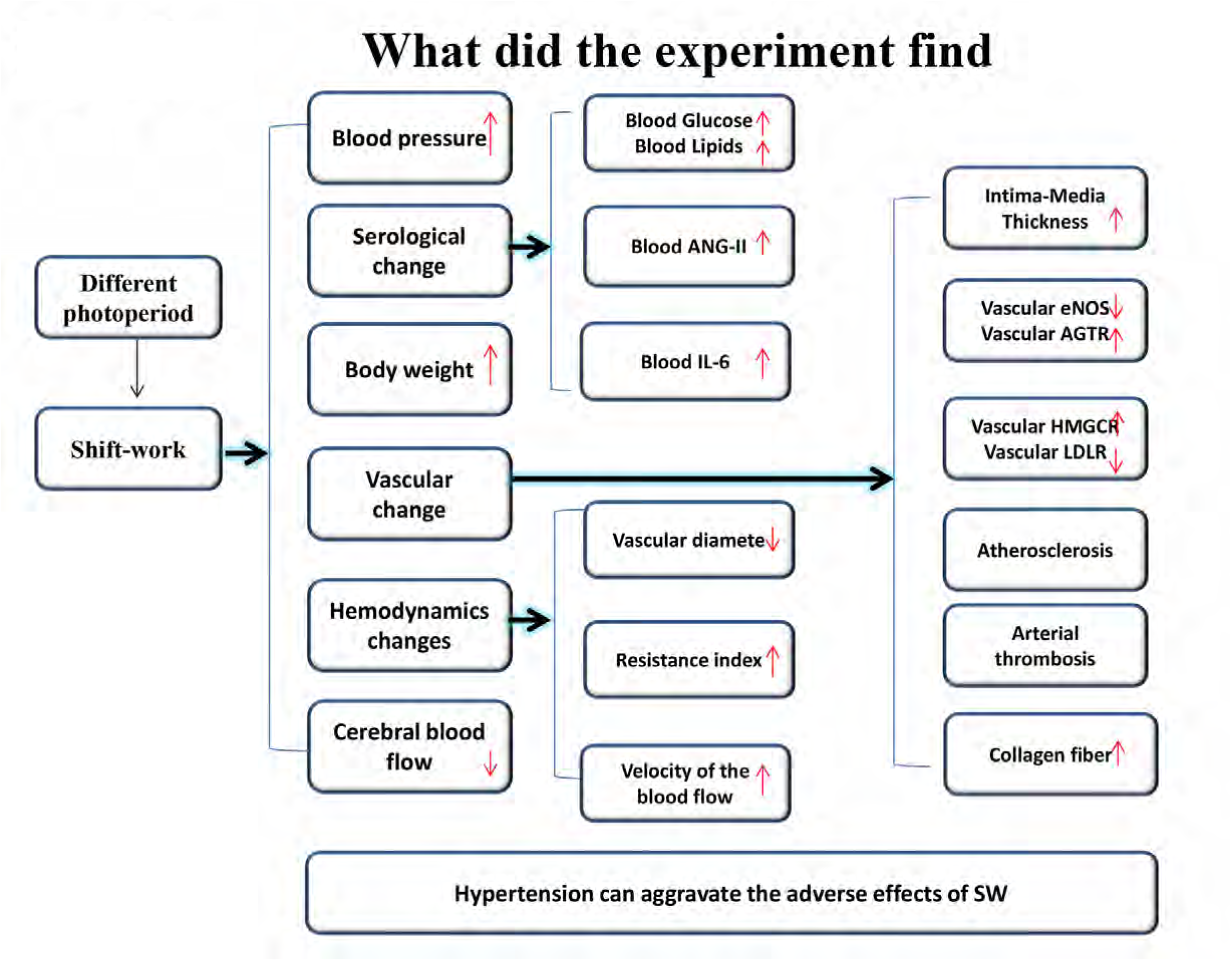
Series of experimental results found in this experiment.

The present study also found that vascular lesions, changes in blood biochemistry, effects on blood pressure, and extent of thrombosis caused by the different types of SW differed. The ClockLab results showed that the rest time in the ILD16:8 group was longer than that for the ILD12:12 group; there were no differences in the other behavioral test results. Although the difference between the two SW groups was not significant in the serological examination, the mean levels of IL-6, ANG-II, APOE, and Glu in the ILD12:12 group were higher than those measured in the ILD16:8 group. Although there was no significant difference in the vascular wall measurements, the rate of carotid arterial thrombosis, intima-media thickness, and collagen fiber ratio in the ILD12:12 group were higher than those recorded in the ILD16:8 group. This indicates that rats in a SW environment with short periods of rest are exposed to a higher risk.

In addition, the present study found differences in the experimental results between the WKY rats and SHR. The two types of rats were classified into isogenic control groups, and their main physiological difference is that the blood pressure of the SHR was significantly higher than that measured in the WKY rats. With regard to behavior, the SHR were more active than the WKY rats [^90, 91^]. These differences in physiological and behavioral characteristics may be responsible for the observed differences in the experimental results between these two types of rats. In the ClockLab behavioral analysis of the number of running wheel revolutions, the SHR were confirmed to be significantly more active than the WKY rats (Figure 3D). The serological examination showed that the levels of TG in the serum of the SHR were lower than those reported in the WKY rats, and that the expression of HMGCR in the vascular wall was higher in the former than in the latter. This suggests that an increase in the amount of exercise may induce beneficial changes in lipid metabolism. Furthermore, due to the hypertension factors in the SHR, the effects of SW on the carotid arteries of these rats were significantly more pronounced than those observed in the WKY rats. Firstly, regarding the carotid arterial anatomy, the intima-media thickness of the carotid artery in the SHR was significantly greater than that recorded in the WKY rats. In addition, the levels of ANG-II in the serum and expression of AGTR in the vascular wall of the SHR were higher than those measured in the WKY rats. These findings indicated that the effects of SW on the decrease in carotid arterial compliance in the SHR were more significant. Secondly, concerning the hemodynamics, CBF was more pronounced in the SHR (SW) than in the WKY (SW) rats. Therefore, the overall effect of SW on the carotid artery of the SHR was significantly greater than that observed in the WKY rats. Thirdly, SW increased the mean arterial pressure in the WKY rats and SHR. Interestingly, SW exerts different effects on both types of rats. In the WKY rats, SW increased the diastolic blood pressure; however, it did not exert a significant effect on the systolic blood pressure. In SHR, ILD12:12 increased the systolic and diastolic blood pressure in SHR, while ILD16:8 increased the systolic blood pressure without significant changes in the diastolic blood pressure. This suggests that changes in blood pressure may differ between normotensive and hypertensive patients when subjected to SW. In fact, the pathological changes and treatment options for systolic hypertension and diastolic hypertension are not identical [^92, 93^]. Thus, further study is warranted to determine the effect of SW on blood pressure.

The present study had several limitations. Firstly, it did not investigate the factors related to arterial thrombosis (e.g., coagulation factors and fibrin). Secondly, the sample size in this study was small. Thirdly, the measurements in the ILD12:12 and ILD16:8 groups did not differ significantly; however, the mean values differed. This indicates that the observation time of 3 months may be insufficient, and the time of observation should be extended accordingly.

In summary, the present study confirmed the that SW may: (1) cause intima-media thickening of the carotid artery and decrease elasticity, compliance, and CBF, and be a risk factor of metabolic disorders; (2) induce atherosclerotic lesions in the abdominal aorta; (3) cause carotid arterial thrombosis by inducing metabolic disorders; and (4) affect CBF by interfering with the anatomy and physiological function of the carotid artery.(5) Hypertension can aggravate the adverse effects of SW.

## Experimental materials and methods

The data of this study are available from the corresponding author upon reasonable request. Details of the major resources may be found in the online-only Data Supplement.

### 1. Experimental animals and groups

Based on the pre-experimental results combined with the statistical calculations of the sample size, six groups were formed (nine rats per group) involving a total of 54 rats. Overall, 27 adult male SHR rats and 27 isogenic, age- and weight-matched WKY rats were included (age: 10 weeks). The R language (version 3.5.0) was used to randomly divide the two types of rats into six groups: WKY (LD12:12), WKY (ILD16:8), WKY (ILD12:12), SHR (LD12:12), SHR (ILD16:8), and SHR (ILD12:12). Subsequently, the blood pressure of the 54 rats was measured, and a statistical analysis was performed.

All rats were purchased from Capital Medical University (Beijing, China) and housed in the Capital Medical University specific-pathogen-free barrier animal facility (Beijing, China). The animal facility was illuminated by 100-W fluorescent lamps with a light intensity of 150–200 lux, and maintained at an indoor temperature of 20–24°C and air humidity of 40–70%. The rats had *ad libitum* access to food and water. All rat food (Beijing Keao Xieli Feed Co. Ltd., Beijing, China) was conventional “rat maintenance feed” with corn, beans, and flour as the main raw materials, and a fat content of <40 g/kg. The average calorie value of rat food was 3.4 kcal/g.

This study was approved by the Institutional Animal Care and Use Committee of Capital Medical University (AEEI-2017-093). The animal experiments were conducted in accordance with the International Guiding Principles for Biomedical Research Involving Animals established by the World Health Organization.

### 2. Model establishment

All rats were housed under normal LD conditions (LD12:12) for 2 weeks to acclimatize them to the experimental environment. After 2 weeks, the activity rhythm of the rats was adjusted using different LD cycles. The LD cycle conditions for the LD12:12 groups (Figure 1A) were as follows: light: 08:00–20:00 and dark: 20:00– 08:00 the following day. The LD cycle conditions for the ILD12:12 groups (Figure 1B) were as follows: light: 08:00–20:00 and dark: 20:00–08:00 the following day for a total of 3 days, and subsequently changed to dark: 08:00–20:00 and light: 20:00–08:00 the following day for a total of 3 days. The LD cycle conditions for the ILD16:8 groups (Figure 1C) were as follows: light: 08:00–20:00 and dark: 20:00–08:00 the following day for a total of 3 days, and subsequently changed to dark: 12:00–20:00 and light: 20:00–12:00 the following day for a total of 3 days. These three models were cycled for 90 days.

### 3. Materials and methods

#### 3.1. ClockLab behavioral analysis

The basic unit consisted of a rat cage with a running wheel (45 cm × 25 cm × 20 cm). Each unit had a metal running wheel (approximately 12 cm in diameter). Each experimental unit was connected to the ClockLab signal transmitter (ACT-500, USA) for signal-to-data transmission. On day 60 of model establishment, we randomly selected six rats from each group (36 rats in total). Each rat was subjected to behavioral analysis for 12 consecutive days. The 24-h activity rhythm of the rats was recorded using the ClockLab software (ACT-500). After the experiment, the MATLAB (2014)-based ClockLab (2.7.3) toolbox was used to plot the rhythm activity plot of the circadian rhythm for data analysis. The rhythm amplitude and period of each rat over the 12 days were analyzed using the “Chi-squared Periodogram” package. The average rest time and number of running wheel revolutions per day for each rat were analyzed using the “Activity Profile” package.

#### 3.2. Blood pressure measurement

The phase for the SW group changed compared with that for the LD12:12 group. However, the LD cycle condition changed at one cycle every 6 days (Figure 1), and 15 complete cycles were completed over the 90-day period of model establishment. Thus, the phase for the SW group at days 91–93 was consistent with that for the LD12:12 rats. In this study, measurements of blood pressure and body weight, serology, vascular ultrasound and arterial spin labeling (ASL) examinations, and carotid arterial section analysis were performed on days 91–93. Both the LD12:12 and SW groups were evaluated at ZT 1 (09:00) to ensure that all rats were evaluated at the same ZT (Figure 1). Blood pressure was measured on day 91, and both the LD12:12 and SW groups were measured at ZT1 (09:00). All 54 rats were assessed.

Method of blood pressure measurement: Blood pressure was measured using a six-channel rat non-invasive CODA sphygmomanometer (KENT, USA). During the measurement, the rats were placed in a fixed container for 5–10 min. After the emotion was stabilized, the Ocuff and VPR sensors were sequentially placed on the base of the tail, and the tail of the rat was placed on a heating plate. When the blood flow reached the sensor threshold, the operation interface displayed the sensor pressure curve and blood pressure value. Five stable blood pressure measurements were performed, and the systolic blood pressure, diastolic blood pressure, and mean arterial pressure were recorded. The rats were maintained in a relaxed state throughout the procedure.

#### 3.3. Weight measurement

Weight measurement was conducted at ZT1 (09:00) on day 91. The weights of all 54 rats were measured.

#### 3.4. Serology

This test was conducted at ZT 2 (10:00) on day 91. Blood was collected from each group of rats immediately after blood pressure measurement. Six rats were randomly selected from each group, and blood was collected from the tail. Approximately 2–3 mL of blood was collected from each rat. After standing for 2 h, the cells were centrifuged, and the serum supernatant was collected. Enzyme-linked immunosorbent assay (ELISA) was performed on a portion of the serum of each rat, while the other portion was used for serum biochemical testing.

ELISA included apolipoprotein E (APOE) (product lot number: CSB-E09749r), interleukin-6 (IL-6) (product lot number: ERC003.96), and angiotensin-II (ANG-II) (product lot number: CSB-E04494r). ELISA was performed using the serum of all six rats from each group. The antibody was first diluted to a protein content of 1–10 μg/mL using carbonate coating buffer. Thereafter, 200 µL of blocking solution was added to each well, and the cells were washed after incubation at 37°C for 1–2 h. The sample was added after washing. After sealing with a film, the plate was incubated at 37°C for 1–2 h and re-washed. Following the addition of the antibody, the plate was re-incubated at 37°C for 1 h and re-washed. A pre-diluted enzyme conjugate working solution was added to each well, and the samples were re-washed after incubation for 30 min. After addition of the chromogenic substrate, the reaction was terminated, and the results were determined.

The serum biochemical tests included determination of the levels of serum glucose (Glu), total cholesterol (TC), triglycerides (TG), high-density lipoprotein (HDL), and low-density lipoprotein (LDL). The serum biochemical analyzer was obtained from Shenzhen Rayto Life and Analytical Sciences Co., Ltd. (product model number: Chemray 240). Serum samples were placed in the analyzer, and the concentration values were calculated. Six rats from each group were selected for the serum biochemical tests.

#### 3.5. Vascular ultrasound

Vascular ultrasound examination was performed at ZT 3 (11:00) on day 91. Carotid artery ultrasound was performed immediately after completion of blood collection. Six rats were randomly selected from each group. Vascular ultrasound measurements of the carotid artery were performed bilaterally using a high-resolution small-animal color Doppler ultrasound imaging system (Vevo 2100; FUJIFILM VisualSonics, Inc.). Vascular ultrasound measurements included the bilateral common carotid arteries, internal carotid arteries, and external carotid arteries. M-mode ultrasound scanning uses multiple scans to observe the anatomy of the blood vessels. The measurement point for the common carotid artery was located approximately 3–4 mm below the bifurcation into the internal and external carotid arteries. The measurement point for the internal carotid artery was located approximately 0.5–1 mm above the bifurcation. Measurements included the systolic luminal diameter, diastolic luminal diameter, intima-media thickness, and presence or absence of plaque formation. Following the detection of plaque formation, the number and area of the plaques were recorded. Hemodynamic measurements were performed using color Doppler ultrasound. The probe was placed in the center of the lumen at the measurement point. The mean velocity of the blood flow and the resistance index were recorded by manually adjusting the angle between the direction of the sound beam and the direction of blood flow to <60°.

#### 3.6. Hippocampal plane ASL examination

This examination was performed at ZT1 (09:00) on day 92. Five of the six rats from each group that had undergone vascular ultrasound examination were randomly selected for magnetic resonance scanning (total: 30 rats). Magnetic resonance scanning of the rats was performed using a 7.0 T magnetic resonance scanner (PharmaScan; Bruker Biospin, Rheinstetten, Germany) at Capital Medical University. The ASL sequence was selected from the hippocampal plane with a slice thickness of 2 mm, field of vision of 30 × 30 mm, echo time of 25 ms, repetition time of 18,000 ms, and fractional anisotropy of 90°. The full thickness of the hippocampal plane of the rats was manually selected, and the CBF was recorded.

#### 3.7. Arterial section analysis

On day 93, the rats were sacrificed at ZT1 (09:00), and the bilateral carotid arteries and aorta were harvested. Samples were obtained from the five rats in each group that had undergone ASL sequence examination (total: 30 rats in the six groups).

Experimental method: The rats were anesthetized via intraperitoneal injection of 1% sodium pentobarbital. The ribs were cut using surgical scissors, and the thoracic cavity was exposed. A perfusion syringe was used to penetrate the left ventricle of the heart; the perfusion syringe was fixed; and 500 mL of normal saline was perfused into the blood vessels. Subsequently, fixation by perfusion with 200 mL of 4% paraformaldehyde was performed. The bilateral carotid arteries and aorta were removed and placed in a paraformaldehyde solution. Hematoxylin-eosin (HE) staining, Masson’s trichrome staining, and immunohistochemistry specimens of the carotid arteries were dehydrated after 48 h of paraformaldehyde fixation and embedded in paraffin to form paraffin blocks. In addition, part of the specimen of the bifurcation of the internal and external carotid arteries was collected to produce frozen sections for Oil Red O staining. The site for common carotid artery specimen collection was located approximately 2–4 mm below the bifurcation of the internal and external carotid arteries. The sites for internal and external carotid artery specimen collection were located approximately 0.5–1 mm above the bifurcation. The aorta was examined via gross Oil Red O staining.

Carotid artery Oil Red O staining: According to the literature, carotid arterial atherosclerosis is more common at the bifurcation of the internal and external carotid arteries. Therefore, frozen sections of specimens collected from the carotid artery bifurcation were used for staining. Initially, the frozen sections were fixed. After fixation, Oil Red O staining was performed, followed by background differentiation. Following differentiation, hematoxylin counterstaining was performed, and the slides were finally mounted. After completion of staining, the proportion of Oil Red O-positive blood vessels and the area of atherosclerosis were calculated.

Carotid artery HE staining: Paraffin sections of the specimens collected from the carotid artery bifurcation and the common carotid artery were used. The paraffin sections were deparaffinized with water and stained with hematoxylin. Eosin staining was subsequently performed. Finally, the slides were dehydrated and mounted. After staining, the intactness and organization of the nuclei of the vascular wall were determined. In addition, the formation of solid plaque in the blood vessels was also examined.

Immunohistochemistry: Changes in the expression of the ANG receptor (AGTR) (product lot number: BA0582), nitric oxide synthase (eNOS) (product lot number: bs-0163R), LDL receptor (LDLR) (10785-1-AP), and 3-hydroxy-3-methylglutaryl-CoA reductase (HMGCR) (13533-1-AP) in the vascular wall were observed via immunohistochemistry. Paraffin sections of the common carotid artery were collected for immunohistochemical analysis. Firstly, the slides were placed in citrate antigen retrieval buffer (pH, 6.0) for antigen retrieval. Secondly, one drop of 3% hydrogen peroxide was added to each slide, and peroxidase activity was blocked via incubation for 25 min at a room temperature (20–24°C). Thirdly, 3% bovine serum albumin was added dropwise to uniformly cover the tissue, and the cells were blocked for 30 min at room temperature. The primary antibody was added dropwise and incubated overnight at 4°C. The secondary antibody was added dropwise and incubated for 50 min at room temperature. Finally, diaminobenzidine color substrate was added dropwise; the nuclei were counterstained, and the slides were dehydrated and mounted. After staining, the total densities of the target proteins were calculated.

The aforementioned sections were obtained using the TissueGnostics multispectral quantitative analysis system. Image acquisition was performed using a (40×) high-power microscope. Image analysis of the sections was performed using the ImageJ software.

Gross Oil Red O staining: Forceps were used to remove the adipose tissue around the blood vessels. The blood vessels were fixed in a fixative for >24 h. The tissue was removed from the fixative solution and washed twice with phosphate-buffered saline solution. The aorta was cut longitudinally using anatomical scissors, immersed in Oil Red O staining solution for 60 min at 37°C. The specimen was differentiated with 75% ethanol until the fat plaques in the lumen were orange-red or bright red, washed twice with distilled water, and imaged. The area ratio of positive plaque in Oil Red O-positive vessels was calculated. Positive plaque area ratio = positive plaque area / total aorta area.

### 4. Statistical analysis and experimental results

The SPSS 24.0 software was used for the statistical analysis. Statistical plotting and data visualization were performed using GraphPad 7.0. Prior to the statistical analysis, the data were tested for normality. Normally distributed data were expressed as X ± S. For non-normally distributed data, the rank-sum test was used, and the data were expressed as medians (interquartile ranges). A two-factor design was used for the statistical analysis of comparisons between two groups. The *post-hoc* Bonferroni test was used for pairwise comparisons between multiple groups. Owing to the inconsistency in the changes in blood flow velocity between the WKY rats and SHR, one-way analysis of variance (ANOVA) was used for comparison between the groups for the vascular ultrasound hemodynamic examination. The *post-hoc* Bonferroni test was used for pairwise comparisons among multiple groups. A *p*<0.05 denoted statistically significant difference. The paired t-test was used to compare the blood pressure before and after the experiment. Fisher’s exact test was used to compare the rates of atherosclerotic solid plaque formation and carotid arterial thrombosis in the HE slices. For the comparison of carotid arterial sections, the average of both sides was used in the statistical analysis.

## Acknowledgments

We are grateful to the China Rehabilitation Science Institute and the Capital Medical University for their support of this experiment. We thank Qing Xu for her assistance in operating rat ultrasound. We also thank Cramer Steven C for his suggestions.

## Author contributions

YunLei Wang and Tong Zhang designed the experiments. Yan Yu, Fan Bai, YuGe Zhang and YaFei Chi improved experimental equipment. HaoJie Zhang and Shan Gao provided suggestions. YunLei Wang and Tong Zhang wrote this manuscript.

## Nonstandard Abbreviations and Acronyms

SW: shift-work
LD: light:dark
APOE: apolipoprotein E
IL-6: interleukin-6
ANG-II: angiotensin-II
Glu: serum glucose
TC: total cholesterol
TG: triglyceride
HDL: high-density lipoprotein
LDL: low-density lipoprotein
ASL: arterial spin labeling
AGTR: ANG receptor
eNOS: nitric oxide synthase
LDLR: LDL receptor
HMGCR: 3-hydroxy-3-methylglutaryl-CoA reductase
SHR: spontaneously hypertensive rats
WKY: Wistar-Kyoto
ZT: Zeitgeber

